# Unravelling complex hybrid and polyploid evolutionary relationships using phylogenetic placement of paralogs from target enrichment data

**DOI:** 10.1101/2024.06.28.601132

**Authors:** Nora Walden, Christiane Kiefer, Marcus A. Koch

**Author notes:** Corresponding authors:* Nora Walden, Marcus Koch.

## Abstract

Phylogenomic datasets comprising hundreds of genes have become the standard for plant systematics and phylogenetics. However, large scale phylogenomic studies often exclude polyploids and hybrids due to the challenges in assessing paralog status of targeted loci and incorporating them into tree reconstruction methods. Using a target enrichment dataset of 1081 genes from 452 samples from the Brassicaceae tribe Arabideae, including many hybrid and high ploidy taxa, we developed a novel approach to disentangle the evolutionary history of this phylogenetically and taxonomically challenging clade. Our approach extends beyond commonly used gene tree-species tree reconciliation techniques by using phylogenetic placement, a method adopted from metagenomics, of paralogous sequences into a diploid tree. We call this approach Paralog PhyloGenomics (PPG), and show how it allows for the simultaneous assessment of the origins of ancient and recent hybrids and autopolyploids, and the detection of nested polyploidization events. Additionally, we demonstrate how synonymous substitution rates provide further evidence for the mode of polyploidization, specifically to distinguish between allo- and autopolyploidization, and to identify hybridization events involving a ghost lineage. Our approach will be a valuable addition to phylogenomic methods available for the study of polyploids.

## Introduction

Repeated cycles of polyploidization and diploidization in many plant lineages are considered a driving force behind the formation of new species (Soltis et al. 2009; Dodsworth et al. 2016), enhancing diversity and disparity (Clark and Donoghue 2018; Stull et al. 2021), and ultimately resulting in adaptation, radiation and increased diversification rates (Soltis and Soltis 2016; Smith et al. 2018). The discovery of ancient whole-genome duplications (WGDs) in plants began with the detailed analysis of the *Arabidopsis thaliana* genome (Simillion et al. 2002; Bowers et al. 2003). Over the next twenty years a multitude of lineage-specific WGDs across the green tree of life have been identified (Soltis and Soltis 2021; Qiao et al. 2022), typically discovered through sequencing entire genomes or transcriptomes (e.g. One Thousand Plant Transcriptomes Initiative 2019). Such WGDs may result from genome doubling within a single taxon (autopolyploidy) or hybridization between different taxa (allopolyploidy) not necessarily representing closely related taxa (Burns et al. 2021), eventually combining different evolutionary lineages (German and Friesen 2014; Mandáková et al. 2018; Walden et al. 2020a). Determining the mode of polyploidization is challenging, particularly for WGDs that occurred millions of years ago (Thomas et al. 2017; Zwaenepoel and Van de Peer 2019). Even for known and more recent hybrid origins, such as in allohexaploid *Brassica* (Chalhoub et al. 2014), phylogenetic trees often fail to accurately depict species’ evolutionary histories. This is because processes like hybridization and incomplete lineage sorting cannot be easily represented in simple bifurcating trees (Huson and Bryant 2006).

Even when utilizing large phylogenomic datasets phylogenetic reconstruction often relies on the use of orthologs; gene sequences that originated, segregated and were sorted through speciation rather than gene duplication (Fitch 1970; Koonin 2005). Orthology can be inferred through synteny (Walden and Schranz 2023) if whole-genome sequences are available, but is more challenging to detect in most phylogenomic datasets that lack complete genomes and detailed synteny information. Consequently, target enrichment studies frequently filter for single-copy loci either during the probe design phase (Johnson et al. 2019; Walden and Schranz 2023) or after sequencing (Jin et al. 2023; Scheunert et al. 2023).

However, orthology is not guaranteed with single-copy loci. Firstly, differential gene loss (fractionation) following WGD can result in the inclusion of pseudoorthologs with single-copy genes (Xiong et al. 2022; Fedosov et al. 2024). Secondly, the differential assembly of paralogous contigs, (gene) sequences that originated through duplication instead of speciation, can create artefactual orthologs (Frost et al. 2024), particularly when data quality and sequencing depth of target enrichment studies are insufficient to assemble full length contigs of all paralogs. Thirdly, orthology inference based on sequence similarity may be biased by user-specified input parameters that cluster sequences into orthogroups. Additionally, single-copy gene sets often contain genes with conserved nucleotide sequences and gene functions that are under selection to revert to single-copy status after duplication (De Smet et al. 2013; Li et al. 2016; Walden and Schranz 2023) In allopolyploids, genes from one of the parental genomes are often preferentially retained, referred to as subgenome dominance (Cheng et al. 2018; Emery et al. 2018), introducing another bias that may dilute the signal of hybridization recoverable in phylogenomic datasets.

On the other hand, phylogenomic datasets, especially those employing data complexity reduction methods like gene capture experiments, are powerful tools to analyze very large datasets (Zimmer and Wen 2015; Bartoš et al. 2023). The most recent angiosperm phylogenetic analysis (Zuntini et al. 2024) utilized the Angiosperms353 bait set (Johnson et al. 2019), capturing 353 nuclear single-copy genes and including 9506 species from 7923 genera across all angiosperm families. Key findings from this study highlighted the role of WGD and incomplete lineage sorting, revealing that early angiosperm evolution at the order level is characterized by significant gene tree conflicts (Zuntini et al. 2024). Therefore, there remains a crucial need to unravel the phylogenetic information of duplicated genes and integrate them into phylogenetic trees. This integration is essential for reconstructing resolved species trees that reflect the comprehensive evolutionary history of species, encompassing lineage divergence, gene duplication, and hybridization events.

Paralogs can provide valuable information for phylogenetic reconstruction (Zhang et al. 2020; Ufimov et al. 2022). Recent studies have shown that the use of larger gene families can enhance phylogenetic signal (Smith et al. 2022). Additionally, gene or genome duplications can be detected, and their position on a species tree can be pinpointed using gene tree-species tree reconciliation methods (e.g. Rabier et al. 2014; Smith et al. 2015; Thomas et al. 2017; Paszek and Górecki 2018). However, the reliability of WGD detection through these methods remains uncertain, and a significant limitation is the requirement of an *a priori* species tree (Zwaenepoel et al. 2019). Methods for obtaining and/or phasing paralog sequences from target enrichment data (Johnson et al. 2016; Nauheimer et al. 2021; Ufimov et al. 2022; Freyman et al. 2023) and species tree reconciliation methods that accommodate the presence of paralogs (Zhang et al. 2020) have become widely used. These approaches hold promise for improving the resolution of phylogenetic trees by integrating the complexities introduced by gene duplications.

While some methods to disentangle paralogs explicitly work for allopolyploids and are even able to phase gene copies according to their parental lineage (Freyman et al. 2023), they typically require prior knowledge of past hybridization events and a resolved species tree. These methods are often limited to small datasets with one or few well-defined hybridization events. This may in part also be due to the limitations posed on phylogenetic reconstruction caused by the reliance on bifurcating trees for large datasets, in which hybrid taxa cannot be properly displayed. Reticulate network analyses of phylogenomic datasets can account for hybridization between taxa (Solís-Lemus et al. 2017) and even extend the methodology to polyploids (Yan et al. 2022). However, computational limitations restrict such analyses to very small datasets (20-30 samples) and few hybridization events (Solís-Lemus and Ané 2016; Jin et al. 2023). Methods from population genetics to identify levels of introgression, such as D-statistics and fbranch analysis (Malinsky et al. 2021), have also been combined with species tree reconstruction and network analyses in smaller phylogenomic datasets such as in *Osmanthus* (Li et al. 2023) and *Alchemilla* (Morales-Briones et al. 2021). Nonetheless, these approaches still rely on a relatively resolved species tree and may be restricted to subsets of data.

Sequencing costs have decreased substantially in recent years leading to the availability of larger and more complete datasets for plant systematics. However, appropriate analysis methods that can account for complex evolutionary patterns are still lacking. Highly polyploid taxa, which may have undergone multiple, nested WGDs are rarely analyzed in their full complexity. Instead, they are often simplified through the use of strict single-copy loci or consensus sequences in phylogenetic studies. An additional confounding factor is that the ploidy status of many plant species remains unknown and cannot easily be determined using available herbarium vouchers, which may be the only material accessible for study.

To address these challenges, we introduce a novel approach that incorporates paralogs assembled from target enrichment data into the reconstruction of an evolutionary hypothesis using phylogenetic placement, a method commonly used in metagenomics (Czech et al. 2022). This method requires no prior knowledge of hybridization, WGD or ploidy status. We apply this approach to a dataset of 452 samples from the Brassicaceae tribe Arabideae as a model system. Arabideae is the largest tribe within Brassicaceae, containing approximately 550 of the family’s 4000 species, including the largest genus (*Draba*) with ∼400 species (Karl and Koch 2013). The tribe belongs to the recently defined subfamily Brassicoideae, supertribe Arabodae (German et al. 2023).

*Arabis alpina*, the taxonomic type of the tribe, has served as a model system for studying annual/perennial lifestyle (Kiefer et al. 2017; Wötzel et al. 2022) and flowering time (Madrid et al. 2021). Several genomes from the tribe have been sequenced in recent years, providing valuable models in comparative genomics (Willing et al. 2015; Nowak et al. 2021). Despite the availability of large phylogenomic datasets, the phylogenetic position of the tribe within the family is debated. Nuclear-based phylogenetic reconstructions (Kiefer et al. 2019; Wötzel et al. 2022; Hendriks et al. 2023) and plastome based reconstructions (Walden et al. 2020a; Hendriks et al. 2023) suggest different positions, while their genome structure shows some affinity to the first diverging tribe Aethionemeae (Walden et al. 2020b). The Arabideae contain a high number of (neo)polyploid species, about 63%, compared to approximately 43% across the entire Brassicaceae (Hohmann et al. 2015), making it taxonomically challenging (Karl and Koch 2013)). However, most major clades within Arabideae have been extensively studied in the past, providing solid background information on taxonomy, biogeography and evolutionary history (Koch et al. 2006, 2010, 2012, 2017; Jordon-Thaden et al. 2010; Karl et al. 2012; Karl and Koch 2013, 2014). This research highlights substantial hybridization, reticulation, and introgression on various spatiotemporal scales, from closely related species and time-scales of few hundred thousand years (Koch et al. 2010; Koch and Grosser 2017) to intercontinental merging of gene pools giving rise to new evolutionary lineages millions of years ago in the giant genus *Draba* (Jordon-Thaden et al. 2010).

Given the high prevalence of polyploidy and the potentially complex history of lineage divergence and hybridization, this Arabideae dataset is an ideal candidate to test our new Paralog PhyloGenomics approach to unravel evolutionary histories using paralogs obtained through gene target enrichment strategies.

## Materials & Methods

### Taxon sampling

Plant material from across the model study system tribe Arabideae (supertribe Arabodae) and outgroup taxa from tribes Boechereae, Turritideae (both supertribe Camelinodae) and Fourraeeae (supertribe Brassicodae) was collected from herbarium specimens, silica dried leaf material and DNA stocks from samples included in previous studies (Supplementary Table S1). 149 of the samples utilized have been investigated in other studies (Koch and Al-Shehbaz 2002; Jordon-Thaden et al. 2010; Koch et al. 2010, 2017; Karl et al. 2012; Karl and Koch 2013; Kiefer et al. 2017) and DNA samples were available. In total we included 452 samples representing 387 accepted taxa (*Arabis* 68 samples, *Draba* 271 samples, *Aubrieta* 18 samples and 29 other samples), which included also five samples from our previous Brassicaceae-wide study (Hendriks et al. 2023). Further information on specimen and vouchers is available in Supplementary Table S1.

Chromosome counts for 179 Arabideae taxa from our dataset were extracted from BrassiBase (Kiefer et al. 2014), and data from an additional 19 taxa was added from PloiDB (Halabi et al. 2023). Base chromosome number in the tribe is eight (Hohmann et al. 2015), though many published chromosome counts are not multiples of eight, suggesting descending dysploidy to five to seven chromosomes (Mandáková et al. 2020). We classified chromosome number into three classes (‘eight’, ‘not eight’, and ‘both eight and not eight’ for taxa with different reported chromosome numbers) and ploidy into eight classes (‘diploid’, ‘tetraploid’, ‘hexaploid’, ‘octoploid’, ‘dodecaploid’, ‘hexadecaploid’, ‘low’, and ‘high’; where low and high were assigned to taxa with mixed ploidy levels with average ploidy below and above tetraploid, respectively).

### DNA isolation, target enrichment, library preparation and sequencing

The samples obtained from herbarium specimens were ground using a Precellys24 homogenizer (PEQLAB Biotechnology GmbH, Erlangen, Germany). DNA was then extracted using the Invisorb Spin Plant Mini Kit (Invitek-Molecular GmbH, Berlin, Germany) according to the manufacturer’s protocol. DNA was eluted in 50 µl elution buffer. DNA concentration was measured by Qubit (dsDNA HS Assay Kit, Invitrogen/Thermo Fisher Scientific, Waltham, Massachusetts, USA) and DNA integrity was assessed by electrophoresis (1% TAE agarose gel).

Prior to library preparation, the integrity of DNA was checked again on an Agilent Tape Station (Agilent Technologies GmbH, Darmstadt, Germany) using the genomic DNA screen tape and corresponding genomic DNA reagents. Fragmentation was only applied if DNA fragments were > 500 bp. For fragmentation the samples were diluted with water in order to obtain an amount of 200 ng of DNA in a volume of 60 µl. Fragmentation was performed using Covaris micro Tubes AFA Fibre with pre-slit SnapCap (6×16mm; Covaris Ltd, Brighton, UK) in a Covaris ultrasonicator (Covaris Ltd, Brighton, UK) for 80 seconds (high fragmentation) or 60 seconds (low fragmentation).

Libraries were prepared using the NEBNext Ultra II DNA library Kit (New England Biolabs GmbH, Frankfurt, Germany) with purification beads and 100 ng of the fragmented/non-fragmented DNA in a volume of 25 µl were used as input according to the manufacturer’s protocol with some alterations. For the NEBNext End Prep (New England Biolabs GmbH, Frankfurt, Germany) half the amount of chemicals was used. For adapter ligation the adapters were diluted 1:15 and half of the amount stated in the protocol was used. Fragments were size selected based on the beads included in the kit (400 to 500 bp).

For the PCR enrichment of adapter-ligated DNA the forward and reverse primer were already combined and NEBNext Multiplex Oligos (New England Biolabs GmbH, Frankfurt, Germany) for Illumina Set 1 to 3 were used as dual index primer pairs in a PCR with 15 cycles according to the manufacturer’s protocol. The PCR reactions were cleaned up using NEBNext sample purification beads (New England Biolabs GmbH, Frankfurt, Germany). Concentration of the libraries was determined by Qubit using the Qubit dsDNA HS Assay Kit (Invitrogen/Thermo Fisher Scientific, Waltham, Massachusetts, USA) and library size was measured on an Agilent Tape Station using D1000 Screen Tape (Agilent Technologies GmbH, Darmstadt, Germany) and corresponding chemicals.

In a next step, libraries were pooled (32 to 43 libraries; a total of 12 pools). Pool 1 was size selected by E-Gel (Thermo Fisher Scientific, Waltham, Massachusetts, USA). Pools were then purified by using Ampure XP beads (Beckman Coulter Life Sciences, Krefeld, Germany), the concentration was measured again by Qubit dsDNA HS Assay Kit (Invitrogen/Thermo Fisher Scientific, Waltham, Massachusetts, USA), and size of the pools was determined by Agilent Tape Station using Agilent Tapestation D1000 Screen Tape and corresponding chemicals (Agilent Technologies GmbH, Darmstadt, Germany).

For hybridization capture two bait sets (Angiosperms353 and custom bait set for Brassicaceae hereafter called Brassicaceae764, Arbor Sciences) were used as described previously (Hendriks et al. 2021). Baits were mixed 2:1 (Angiosperms353:Brassicaceae764) and hybridization was carried out at 60 °C for 24h in a PCR cycler. Binding to streptavidin-coated magnetic beads and washing of the reactions for removal of non-target DNA was performed according to the manufacturer’s protocol (Arbor Scientific, Arbor Biosciences, Ann Arbor, Michigan, USA). In a last step, libraries were suspended and amplified (NEBUltra Q5 polymerase (New England Biolabs GmbH, Frankfurt, Germany), IDT xGen (IDT, Coralville, Iowa, USA) amplification primer; 18 cycles; all according to manufacturer’s protocol). Enriched pools were purified using Ampure XP beads (Beckman Coulter Life Sciences, Krefeld, Germany). Reamplification of the purified PCR product was performed for 10 cycles using the same polymerase and primer as stated above. Final purification was again done using Ampure XP beads (Beckman Coulter, Krefeld, Germany). Concentration of the pools was determined by Qubit (DNA HS Assay, Invitrogen/Thermo Fisher Scientific, Waltham, Massachusetts, USA) and pools were checked on an Agilent Tape Station (D1000 Screen Tapes, D1000 reagents, Agilent Technologies GmbH, Darmstadt, Germany). 25 µl of each pool were sequenced by Illumina sequencing technology (150bp paired end reads; 48 to 65 GB data per pool) at Novogene UK. All raw reads are available at ENA (https://www.ebi.ac.uk/ena/browser/home) under accession PRJEB72669.

### NGS pipeline

Raw sequencing reads were first processed with Trimmomatic 0.39 (Bolger et al. 2014) to remove remaining adapter sequences and trim reads for quality with leading and trailing quality 20, a quality of 20 in a sliding window of 4 bp and minimum length 30. Sequence-based ploidy inference was conducted on trimmed reads. KMC 3.1.1 and kmc_tools 2.3.0 (Kokot et al. 2017) were used to calculate kmer frequencies and generate a histogram of kmers using kmer length 17 and a maximum count of 10,000. Smudgeplot (Ranallo-Benavidez et al. 2020) was then used to infer ploidy level using a lower coverage limit of 4 and an upper limit estimated from the kmer histogram. Ploidy level information from the literature was available for 198 taxa (corresponding to ∼52% of the accessions in our study), and kmer-based ploidy level estimates were used herein to (1) allow comparisons among datasets and (2) apply ploidy level estimates across the entire dataset.

Our data analysis pipeline for the following steps is illustrated as a flowchart in Figure 1. Assembly of gene sequences was conducted with HybPiper 2.0 (Johnson et al. 2016) using the 1081 target genes as a reference (Johnson et al. 2019; Nikolov et al. 2019). As the two enrichment gene sets were developed independently, some genes were contained in both. Following (Hendriks et al. 2023), these 36 genes were removed from the Brassicaceae764 set, as the sequences in this set were generally shorter than those of the Angiosperms353 set. We ran the HybPiper command hybpiper assemble in nucleotide mode using bwa for read mapping and subsequently obtained assembly statistics with hybpiper stats. Due to the high mapping rate (Fig. S1), we could apply a strict quality threshold and only select samples with at least 900 genes covered (GenesWithContigs ≥ 900) and at least 50% of genes covered at 50% length (GenesAt50pct ≥ 226) in HybPiper (Fig. S2) in a first filtering step. In a second step, only genes with less than 20% of missing samples were retained for further analysis (Fig. S3). The final dataset comprised 442 samples and 994 genes. In the case of multiple full-length contigs for a locus, HybPiper identifies main contigs first by coverage, and if multiple contigs of similar coverage are found by similarity to the reference sequence. Main contigs for the selected samples were obtained using hybpiper retrieve_sequences, and paralogs (i.e. all full-length contigs regardless of coverage and similarity to reference sequence) using hybpiper paralog_retriever.

**Figure 1.**
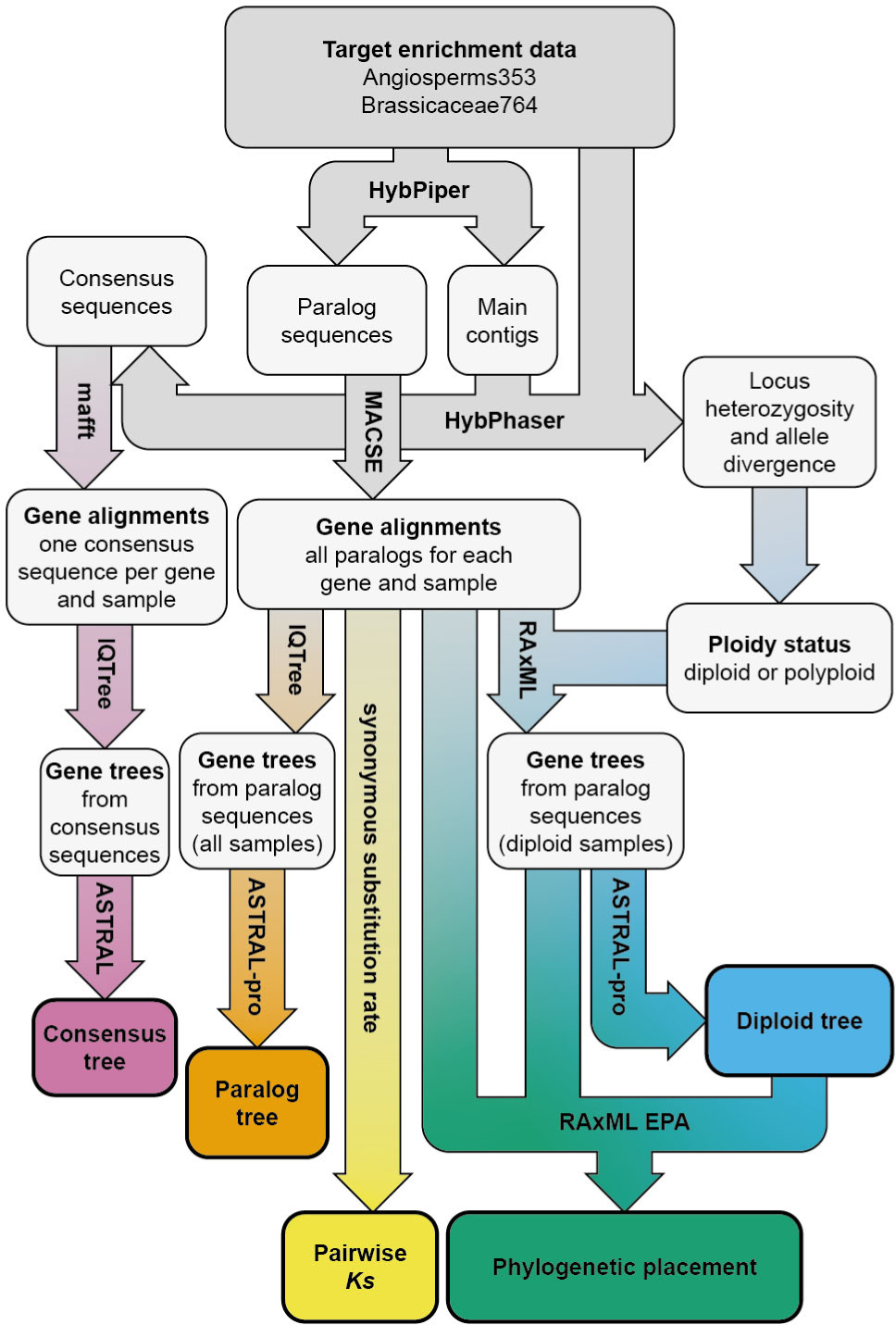
Flowchart of our data analysis pipeline. Target enrichment data (grey) was processed with HybPiper and HybPhaser to obtain consensus sequences, which were aligned and subjected to gene tree reconstruction and finally gene tree-species tree reconciliation to obtain the consensus tree (pink path). Similarly, the paralog tree was reconstructed from gene trees containing paralog sequences obtained from HybPiper (orange path). Pairwise Ks values were obtained from the same alignments (yellow path). Using locus heterozygosity and allele divergence from HybPhaser, ploidy status was inferred to obtain subsets of paralog sequences containing only ‘diploid’ samples; the diploid tree was reconstructed using these alignments (blue path). Finally, evolutionary placement of paralog sequences into the diploid tree was performed using RAxML-EPA (green path).

Next, we used additional software to assess paralogs. HybPhaser 2.0 (Nauheimer et al. 2021) was run to generate consensus sequences and to obtain allele divergence and locus heterozygosity of all remaining samples and genes using a minimum depth for inserting ambiguity codes of 8, minimum allele frequency of 0.1 and minimum count for alleles to be regarded for assigning ambiguity code. In consensus sequences obtained using HybPhaser reads are mapped back to the de novo assembled HybPiper gene contigs and heterozygous sites are represented by ambiguity codes, irrespective of their status as alleles or highly divergent paralogs. A previous genus level phylogenomic study across all tribes in Brassicaceae using the same gene sets (Hendriks et al. 2023) had assessed several different thresholds for proportion of samples and loci covered as well as proportion of SNPs to remove loci with paralogs. When using their thresholds for our data set, the strictest filtering setting resulted in only 34 remaining loci (3.4%) that did not contain high numbers of potential paralogs (Supplementary Table S2). In a dataset with high levels of polyploidy however, even these remaining genes may underly differential gene loss, thus the gene trees may not necessarily reflect the same orthologs consistently across the phylogeny. We therefore had to resort to another strategy for phylogenetic reconstruction.

### Phylogenetic reconstruction

A first tree reconstruction was conducted based on the consensus sequences from HybPhaser (subsequently called ‘consensus tree’, pink path in Fig. 1) with 994 genes from 442 samples using coalescent-based species tree estimation (Mirarab et al. 2014; Zhang et al. 2018)): Gene sequences were aligned using mafft 7.511 (Katoh 2002; Katoh and Standley 2013) with default parameters and the alignment was trimmed in trimAl 1.4rev22 (Capella-Gutiérrez et al. 2009) using the noallgaps option. Gene trees were then reconstructed with IQTree 2.2.0 (Minh et al. 2020) generating 1,000 ultrafast bootstrap replicates (Hoang et al. 2018) with GTR+I+G as substitution model, and used as input for species tree reconstruction using ASTRAL 5.7.8 (Zhang et al. 2018).

Second, we reconstructed a species tree based on all paralogs found for the 994 genes from 442 samples using ASTRAL-pro, which allows for species tree reconstruction despite the presence of paralogs (orange path in Fig. 1). Separate nucleotide sequences of all paralogs without chimeric sequences for each gene obtained by HybPiper (366-1146 sequences per gene, mean number of sequences per gene 520; 916-1719 sequences per sample, mean number of sequences per sample 1170) were aligned using mafft 7.511 (Katoh and Standley 2013) using the FFT-NS-2 method and subsequently realigned with refineAlignment in MACSE 2.06 (Ranwez et al. 2018) to obtain codon-aware nucleotide alignments. Alignments were trimmed with trimAl 1.4.rev22 (Capella-Gutiérrez et al. 2009). Gene trees including all paralogs were reconstructed using IQtree as described above. A species tree (subsequently called ‘paralog tree’, orange path in Fig. 1) was then reconstructed from gene trees with paralogs using ASTRAL-pro 1.13.1.3 (Zhang et al. 2020).

Third, we used a multi-step approach to reconstruct the evolutionary history of tribe Arabideae. We split the dataset into ‘diploid’ and ‘polyploid’ individuals based on allele divergence and locus heterozygosity obtained from HybPhaser (Fig. S4, S5), with ‘diploids’ defined as all individuals with allele divergence < 2 and locus heterozygosity < 90. It is important to note that this classification does not necessarily reflect ploidy level according to chromosome number – a ‘diploid’ ploidy status may be inferred for (recent) autopolyploids due to low heterozygosity, and conversely, a ‘polyploid’ status may be inferred for (recent) homoploid hybrids or for old hybrids that subsequently underwent rediploidization, due to high heterozygosity. Comparison of ploidy status inferred using this method was highly consistent with data from the literature, but less so with ploidy level inferred using kmer (Fig. S6). Because literature data was only available for 246 of 442 samples, we used the HybPhaser-based ploidy status in the following analyses and identified 141 ‘diploid’ samples. Subsequently, sequences from selected diploid individuals were extracted from the initial alignment for separate analysis. The subset gene alignments were also trimmed with trimAl, and maximum likelihood (ML) gene trees were reconstructed with RAxML v8.2.12 (Stamatakis 2014) using the GTRCAT substitution model, with 1000 rapid bootstraps inferences followed by a thorough ML search. As the diploid dataset still contained some paralogs for some genes (number of sequences per gene 112-285, mean number of sequences 145; number of sequences per sample 916-1058, mean number of sequences per sample 1022), we used ASTRAL-pro (Zhang et al. 2020) to combine the gene trees into a species tree (subsequently called ‘diploid tree’, blue path in Fig. 1).

To obtain the phylogenetic position of each paralog, diploid gene trees were first rooted to the two diploid outgroup species from tribe Boechereae using nw_reroot from Newick utilities (Junier and Zdobnov 2010); in cases where one or both of the outgroup samples were missing or had paralogs, trees were rooted manually in figtree 1.4.4 (http://tree.bio.ed.ac.uk/software/figtree/). Phylogenetic placement of paralogs into the diploid gene trees was then conducted using RAxMLs evolutionary placement algorithm (EPA) with the same substitution model as above for each gene (Fig. 1). For allopolyploidy events with known parents, methods to ‘phase’ the two gene copies according to their origin have been described (Freyman et al. 2023). However, in our case the history of WGDs is unknown and could be complex, with multiple and even nested hybridization events between lineages giving rise to entire clades. Using phylogenetic placement of paralogs (green path in Fig. 1) allowed us to pinpoint the placement of the last common ancestor of the extant paralog and the diploid species, and ancient autopolyploidization events as well as hybridization could be inferred from the emerging patterns. RAxMLs EPA outputs the likelihood of placements into each branch as likelihood-weight ratios (lwrs), and we added up the likelihood of all placements of all paralogs across each sample and subsequently normalized branch-specific lwrs by the total sum of lwrs per sample across all 994 genes. This resulted in a unique placement pattern for each ‘polyploid’ sample, though many samples showed similar patterns. We used hierarchical clustering of the normalized placement to group samples with similar patterns and further analyze them in detail. For each cluster, we then considered only placements into branches reaching a mean lwr of at least 0.05 across all samples for visualization in the diploid anchor tree. Filtering of datasets, statistical analysis and plotting of figures was performed in R 4.2.2 (R Core Team 2022) using the following packages: ggplot2 3.4.2 (Wickham 2016), ggpointdensity 0.1.0 (Kremer 2019), viridis 0.6.3 (Garnier et al. 2023), ggpubr 0.6.0 (Kassambara 2023), ggalluvial 0.12.5 (Brunson and Read 2019), ape 5.7-1 (Paradis and Schliep 2019), stringr 1.5.0 (Wickham 2022), treeio 1.22.0 (Wang et al. 2020), ggtree 3.6.2 (Yu et al. 2017), ggExtra 0.10.1 (Attali and Baker 2023), BoSSA 3.7 (Lefeuvre 2020), ggdendro 0.1.23 (Vries and Ripley 2022) and phytools 1.9-16 (Revell 2012).

### Analysis of synonymous substitution rate of paralogs

We further analyzed paralog placement patterns in four clusters as examples for different evolutionary scenarios using pairwise synonymous substitution rates (*Ks*) between and within samples (yellow path in Fig. 1). *Ks* was calculated using the function kaks from R package seqinR (Charif and Lobry 2007). Taxa of interest as well as three outgroup taxa to assess consistency and reliability of the method were analyzed for each cluster and results were displayed with ggridges 0.5.4 (Wilke 2022).

## Results

### Data quality is very high for selected genes and samples

We obtained high read mapping rates for most newly sequenced samples (2.4-82.3%, mean 66.52 %, median 70.7%; see Fig. S1) despite including a number of old herbarium vouchers dating back as far as 1823. This also led to generally high rates of gene recovery; ten samples with fewer than 900 genes with contigs and fewer than half of all genes covered at 50% were excluded from further analysis (Fig. S2). Similarly, the high gene recovery rate allowed us to apply a fairly strict threshold for number of missing samples in each gene (20%) while only excluding 87 genes from downstream analyses (Fig. S3). The final dataset thus consisted of 994 genes from 442 samples.

### Paralogs are prevailing and ploidy is as high as expected

Many selected samples had high numbers of paralogs (5-706 paralogs, mean 159.1, median 140). The highest number of paralogs both with regards to number of genes with paralogs (659) and total sequences (1718) was detected in polyploid *Draba ladyginii*, while *Draba linearifolia* had the lowest number of genes with paralogs (5).

Paralog detection can be influenced by coverage, specifically by the number of reads mapped to the target, as the multiple gene copies are assembled *de novo*. In our dataset, even low numbers (>200,000) of mapped reads were sufficient to identify high numbers of paralogs (Fig. S7a). In line with expectations, we detected significantly lower numbers of paralogs in diploid taxa and those with low ploidy compared to polyploids (Fig. S7b).

Additionally, allele divergence (AD) and locus heterozygosity (LH) were generally high in known polyploid taxa (Fig. S4). We therefore used both measures to identify putative ‘diploids’ based on target sequences rather than literature or raw reads, using a threshold of AD < 2 and LH < 90 (Figs. S5). Samples assigned to the ‘diploids’ group using HybPhaser contained mostly diploid taxa or those with low ploidy levels according to literature, while most tetraploid and hexaploid taxa and all samples with known high ploidy (> hexaploid) were classified as ‘polyploids’ (Fig. S7). Conversely, ploidy assignment based on raw reads using smudgeplot was largely inconsistent with ploidy reported in literature. Interestingly however, most samples from diploid taxa according to the literature which were assigned to ‘polyploids’ using HybPhaser were also inferred to be polyploid using smudgeplot, likely due to their high heterozygosity. With regards to the number of paralogs detected, we found a strong positive correlation with assembly quality (i.e. number of genes assembled in a single contig), suggesting that paralogs may have been missed in ‘polyploid’ samples with lower assembly quality (Fig. S8).

### Phylogenetic reconstruction using conventional methods resolves a simple backbone only

To include the high number of paralogs into phylogenetic reconstruction, we used different approaches: We first reconstructed a species tree from consensus sequences obtained in HybPhaser (pink path in Fig. 1) using all selected genes and samples with ASTRAL (Fig. S9). The major evolutionary clades (*Draba*, *Tomostima* clade, *Aubrieta*, *Drabella*, *Arabis alpina* clade, main *Arabis* clade, *Scapiarabis* clade; Karl and Koch 2013) were generally recovered, though unlike in previous studies *Arabis aucheri* and *Arabis parvula* did not cluster together. Instead, *A. aucheri* was sister to the *Aubrieta* clade and *A. parvula* sister to *Drabella muralis*. Posterior probabilities were high at the backbone, but often low within clades, and quartet scores were low for most branches (Fig. S10).

Next, we reconstructed a species tree from all gene trees based on all paralogs obtained with HybPiper (orange path in Fig. 1) using ASTRAL-pro (Fig. S11). Again, the major clades were recovered, and *A. aucheri* and *A. parvula* formed a clade that was sister to the *Tomostima* clade and *Draba*. However, node support at the respective backbone nodes was low both with regards to posterior probability and quartet score. In contrast to the consensus tree, the quartet score was higher for some nodes in the *Tomostima*, *Aubrieta*, *Drabella* and *Arabis alpina* clades, but lower in *Draba*. The number of quartet trees in all gene trees that supported the first topology was lower when using paralogs (Fig. S12). Both methods were however not able to resolve relationships within the major clades well, in particular in the large genus *Draba*.

### The diploid tree is reliable and suitable for phylogenetic placement

To bypass the potential noise interfering with phylogenetic resolution that may be caused by the high number of paralogs in polyploid samples, we selected 138 ‘diploid’ samples using a maximum allele divergence of 2 and locus heterozygosity of 90% (Fig. S6). Almost all (∼93%) samples with known ploidy in this dataset were either diploid or from a taxon assigned to ‘low’ ploidy level (Fig. S7), indicating a generally high consistency between our detection method for ‘diploids’ and known ploidy levels. However, as sequence divergence between alleles was the prerequisite for ploidy determination, polyploids showing low divergence, such as very recent autopolyploids, may be overlooked.

The diploid tree reconstructed using ASTRAL-pro based on the 994 genes (blue path in Fig. 1) also recovered the major clades of Arabideae (Fig. S13). The *Tomostima* clade as well as *Drabella* and *A parvula* consisted exclusively of ‘polyploids’ and were therefore not included. Using this reduced dataset improved support values; posterior probabilities were largely > 0.95 and more branches had higher quartet scores compared to the consensus and paralog tree (Fig. S14). However, low quartet scores around 1/3 were still found within clades; this may be in part due to a lack of nucleotide differences accumulated during the short time frames in which Arabideae diverged, leading to gene trees with short branch lengths at the respective nodes as well as conflicting topologies.

### Phylogenetic placement into the diploid tree identifies patterns and evolutionary processes

The diploid gene trees and species tree were used as the basis for the next step, phylogenetic placement of paralogs (blue in Fig. 1). Genes with a high proportion (≤10%) of missing samples among diploids were discarded, as were genes with a high paralog proportion (≤10%), resulting in 857 genes (Fig. S15). The pattern of paralog placement likelihood-weight ratios (lwrs) showed clade-specific clustering in particular branches of the tree (Fig. 2, Fig. S16). Note that short internal branches lead to low lwrs at multiple nearby nodes, thus pinpointing the exact branch for each placement was not always possible.

**Figure 2.**
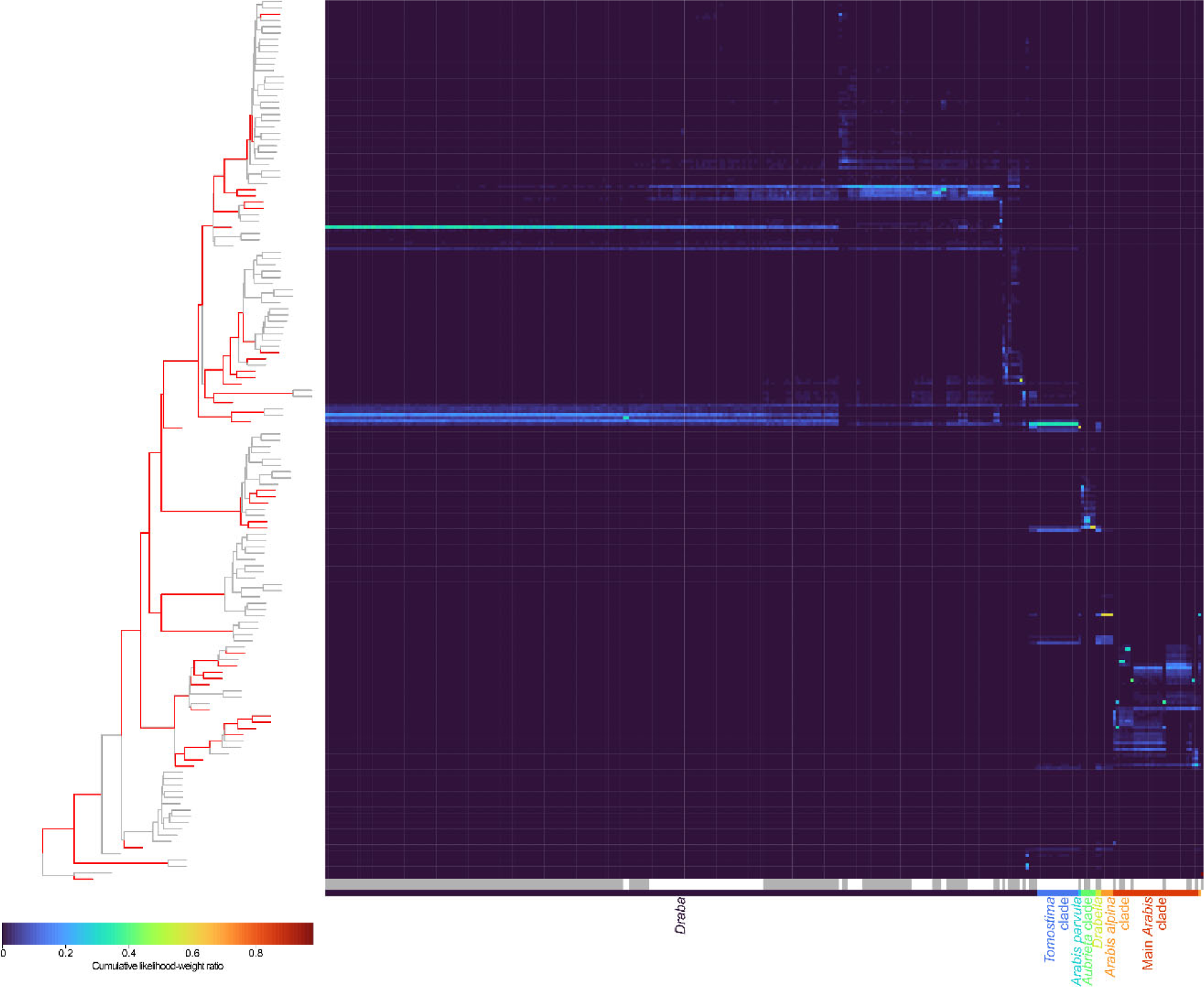
Placement of paralogs. Branches with a cumulative likelihood-weight ratio (lwr) of greater than 0.05 in any cluster are highlighted in red in the diploid tree on the left side; some clades are involved in notably few polyploidization/hybridization events. The heatmap on the right shows lwrs for all polyploid samples (x-axis) across the branches of the diploid tree (y-axis). Similar patterns of lwr across species indicate their shared evolutionary history; the 43 clusters of similar patterns are highlighted in grey and while below the heatmap.

Hierarchical clustering of the placement patterns into 43 clusters (Fig. S17, S18) was used to summarize taxon groups with potentially similar evolutionary histories.

Bringing together the placement patterns and the anchoring diploid tree for each cluster (Fig. S19-S29), we could identify different evolutionary processes based on the number of distinct branches with paralog placement: A majority of paralogs placed into a single branch could indicate a WGD involving only this branch, such as an autopolyploidization event. This pattern was observed in cluster 25 (Fig. S25), cluster 28 (Fig. S25) and cluster 30 (Fig. S26).

Paralogs mapping to two distinct branches indicate ancient hybridization or allopolyploidization (e.g. cluster 1, Fig. S19). Different numbers of paralogs mapping to both involved branches could be a result of asymmetric gene loss after hybridization, or of introgression. Placement patterns consistent with hybridization were found in many clusters in our study, among them clusters 1-3 (Fig. S19) and clusters 31-37 (Fig. S26-S28). The presence of additional WGDs or hybridization events can lead to placement patterns involving more than two distinct branches in the diploid tree, such as in cluster 4 (Fig. S19). This cluster shows a similar placement pattern as clusters 1-3, but with additional paralogs mapping to a third branch, indicating subsequent exchange of genetic material with a third lineage.

Finally, we observed some clusters in which paralogs were placed to many different, neighboring branches along the phylogenetic backbone, for example cluster 29 (Fig. S26). This pattern may be due to a rapid divergence of the major lineages, leading to a lack of nucleotide differences at the targeted genes at the backbone nodes of the Arabideae phylogeny and thus low gene tree resolution. To a lesser extent, this pattern was also observed in other clusters, such as in cluster 30 (Fig. S26), where it could indicate that the inferred WGD occurred shortly after lineage divergence, while in other cases it may simply be indicative of the rapid diversification in the tribe leading to incomplete lineage sorting.

### Synonymous substitution rates and age estimations coincide with evolutionary history

Synonymous substitution rates between paralogs can be used to elaborate on the inferred/hypothesized evolutionary scenarios as well as to provide some relative estimates for the time of polyploidization and hybridization (yellow path in Fig. 1). Here, we chose four exemplary clusters as study cases and analyzed them in more detail using pairwise *Ks* within polyploid samples and between polyploid and diploid outgroups. Clusters representing three different evolutionary scenarios for the origin of hybrids and polyploid taxa (Fig. S30), namely the formation of homoploid or polyploid (allopolyploid) hybrids, autopolyploidization, and hybridization with the involvement of a ghost lineage were chosen.

Example 1 highlights hybridization among two diploid lineages, which hybridized after their divergence and gave rise to a polyploid taxon, which afterwards may undergo further lineage diversification. This is the case in cluster 1 (Fig. 3), where the ancestors of two diploid clades, *Draba polytricha* (as diploid relative A; Fig. 3A) and the clade of *Draba lutescens*, *Draba huetii* and *Draba ellipsoidea* (represented by *D. lutescens* as diploid relative B) gave rise to a large clade of >100 exclusively (according to available data) polyploid taxa (Fig. 2). Paralogs from the polyploid samples (*Draba exunguicula* and *Draba mogollonica*) mapping to the two different branches (Fig. 3B) were analyzed separately here. Within-polyploid *Ks* was similar in both polyploid representatives, confirming their shared evolutionary origin. It was higher than that observed in pairwise comparisons with the closer diploid (Fig. 3C, Fig. S31; closer diploid refers to the taxon that the respective paralog mapped to during the placement analysis) and higher than pairwise *Ks* with the close paralog from the other polyploid, in line with a scenario where two diploid lineages hybridized sometime after their divergence and their hybrid offspring subsequently underwent a rapid lineage diversification.

**Figure 3.**
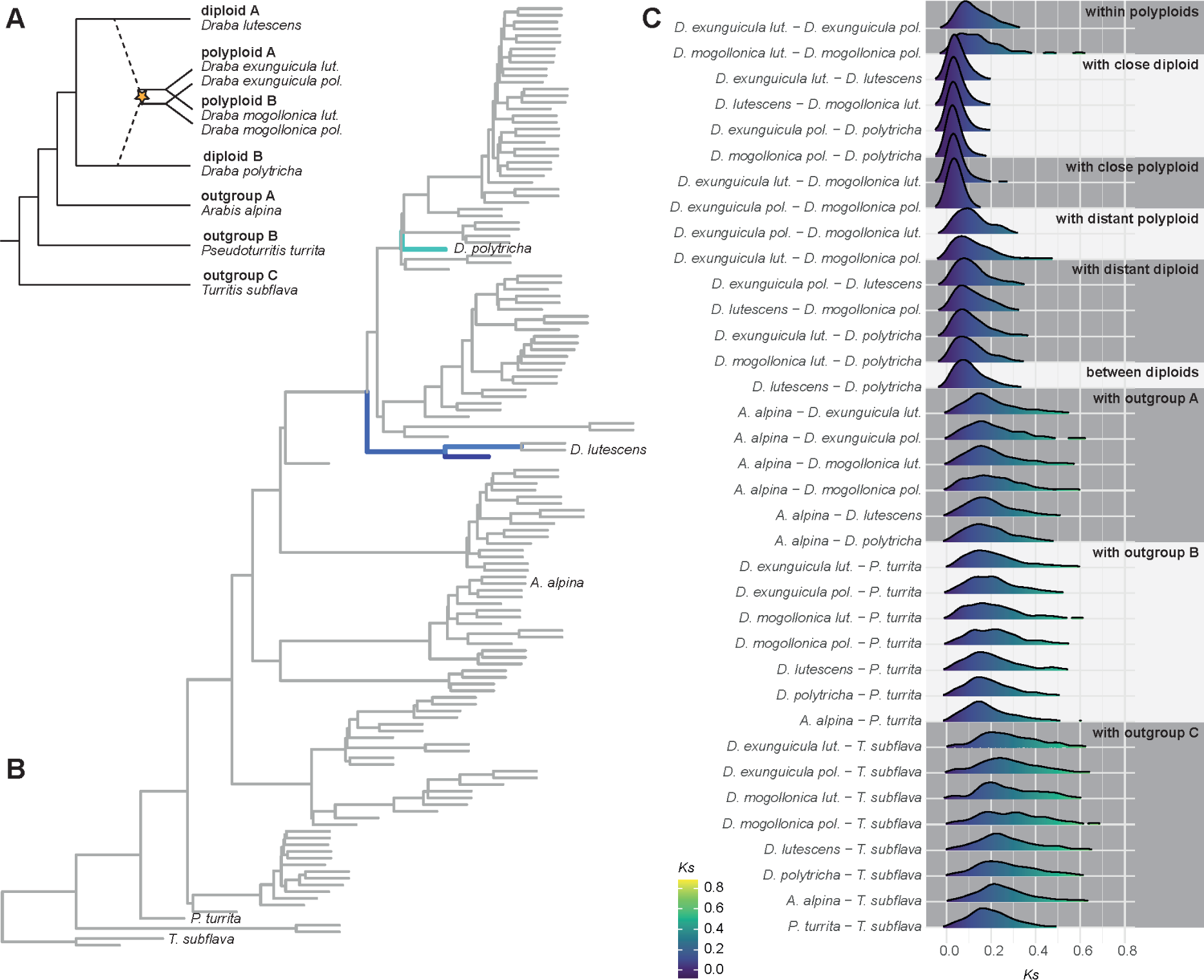
Relative divergence estimation using synonymous substitution rates. Cluster 1 was chosen as an example to illustrate the use of synonymous substitution rates (*Ks*) as a means to confirm the evolutionary scenario inferred from placement patterns and provide a relative timeline. (A) The evolutionary scenario including the hybrid origin of the polyploid clade from two diploid lineages, along with the taxa analyzed in detail. (B) Placement patterns of cluster 1 into the diploid tree. The likelihood-weight ratios (lwrs) at branches with lwr >0.05 are shown, and the taxa selected for detailed analysis of *Ks* are highlighted. (C) Distribution of pairwise *Ks* values between the different species and within polyploids are shown. For polyploids, subsets mapping to the branches leading to diploid A and diploid B, respectively, were analyzed separately.

The second scenario illustrates autopolyploidization, where a single lineage is involved in the emergence of a polyploid clade (Fig. S30). The placement patterns of clusters 25, 28 and 30 suggested such an autopolyploid origin. The polyploid in the second example, cluster 28, was *Arabis verna.* This taxon is sister to the third largest Arabideae genus *Aubrieta* and has been considered a member of the *Aubrieta* clade (Karl and Koch 2013; Koch et al. 2017), which is also in accordance with our results.. Intraspecific *Ks* in *A. verna* was similar to pairwise *Ks* with the two diploid representatives (*Aubrieta canescens* and *Aubrieta macrostyla*) in our analysis (Fig. S32), suggesting a polyploidization event around the time of divergence from the *Aubrieta* lineage that could have involved reproductive isolation driven by ploidy differences.

The third example comes from cluster 30, where three polyploid taxa were placed on a long branch within the *Arabis alpina* clade. Our *Ks* analysis included two of them (*A. mollis* and *A. nordmanniana*) as well as diploid *A. alpina*, and diploid *A. auriculata* from the next sister clade. The first peak in pairwise *Ks* value of *A. mollis* and *A. nordmanniana*, representing the lineage diversification of the polyploid clade, was much lower than that of within-polyploid *Ks* comparisons representing the time of autopolyploidization (Fig. S33), in line with a long post-WGD lag-phase (Schranz et al. 2012). Furthermore, the similar pairwise *Ks* values between polyploids, *A. alpina* and sister clade *A. auriculata* hints at a time of origin shortly after divergence from the *A. auriculata* lineage.

The fourth example assumes involvement of a ghost lineage. The placement pattern of *A. parvula* (cluster 25) indicated a WGD (autopolyploidization). However, in contrast to the two previous examples, *Ks* analysis did not confirm this assessment. Instead, the *Ks* distribution showed an additional peak in the pairwise comparison between *A. parvula* and its closest diploid relative *A. aucheri*, which was considerably lower than intraspecific *Ks* in *A. parvula* (Fig. S34). This pattern can be best explained by the involvement of a ghost lineage: A now-extinct, or unsampled, distant relative of *A. aucheri* originated early and remained isolated for some time, then came into contact with *A. aucheri* more recently, leading to the origin of polyploid *A. parvula* (Fig. S30). Importantly, pairwise *Ks* values with outgroup taxa were consistent with the respective evolutionary scenarios in all four analyzed clusters, and also highly consistent between analyses, indicating their applicability for (relative) time estimates.

## Discussion

The genomics era has produced an enormous and ever-increasing amount of genome scale datasets leading to the discovery of hundreds of recent and ancient whole genome duplications throughout land plant evolution (Vanneste et al. 2014; One Thousand Plant Transcriptomes Initiative 2019; Zuntini et al. 2024). However, current phylogenetic and phylogenomic methods primarily represent evolution as bifurcating trees. The tremendous advances in assembling and analyzing phylogenomic datasets have exposed the limitations of our current methodology to analyze, classify and interpret the evolution of plant clades in the face of ancient hybridization and whole-genome duplication (reviewed in Stull et al. 2023).

This highlights the urgent need for new approaches that can handle more complex evolutionary scenarios. Recently, Smissen and colleagues (2024) developed an innovative approach to infer the evolutionary history of Australasian *Lepidium*, a taxonomically challenging Brassicaceae genus rich in high polyploids, interploidal and intercontinental hybridization. They used amplicon sequencing and condensed splits from MUL-networks. Although their dataset was much smaller than ours, encompassing only 21 samples and 15 genes from a restricted geographical range, their study beautifully illustrates both the demand and the potential for creative solutions to study the origin of allopolyploids in a phylogenetic and systematic context. Similarly, a larger study of the genus *Rosa* used amplicon sequencing and a combination of read-based ploidy estimation, species-tree estimation, and network reconstruction to identify hybrid and polyploid roses and their diploid ancestors (Debray et al. 2022).

Our study system, the Brassicaceae tribe Arabideae, exhibits several of the same challenges commonly faced by current phylogenomic datasets: diverse levels of polyploidy, with ploidy levels ranging up to hexadecaploid, unknown ploidy status in taxa only available as herbarium vouchers, a large evolutionary lineage with diversification into over 500 taxa in under 20 Ma; and an unknown number of (nested) WGD events. In this study, we aimed to develop an approach to investigate the evolutionary history of such a clade using target enrichment data. The outputs at the specific stages of our analysis pipeline are summarized in Figure 4.

**Figure 4.**
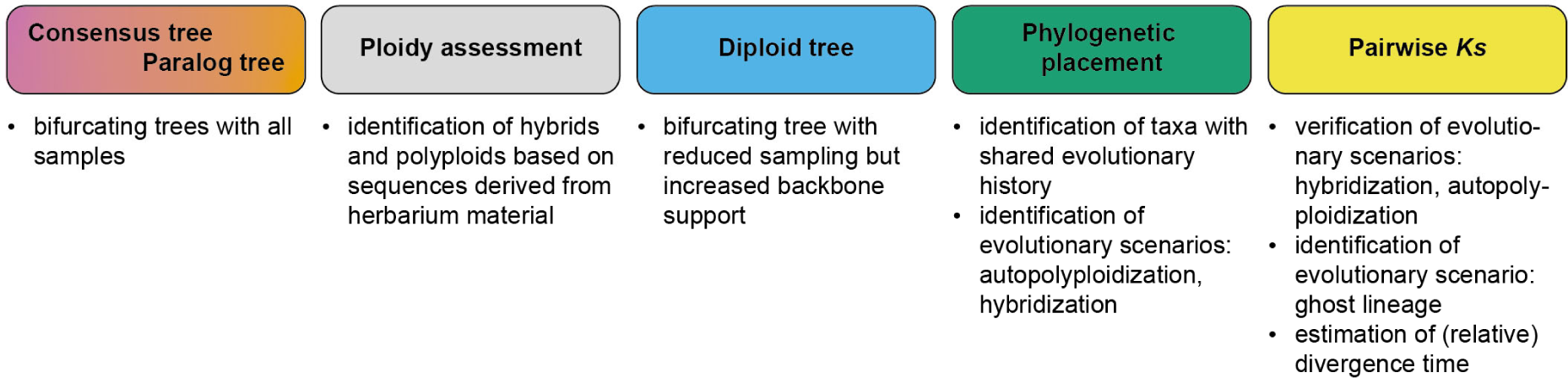
Output summary. The output of the different parts of our analysis pipeline are summarized above. While conventional phylogenomic methods (consensus tree, paralog tree) generate bifurcating trees including all samples, identifying diploids and reconstructing a tree with reduced sampling (diploid tree) provides a phylogeny with increased backbone support. Phylogenetic placement of paralogs furthermore allows us to identify hybrid or polyploid samples with a shared evolutionary history and at the same distinguish between evolutionary scenarios involving autopolyploidization and hybridization. Finally, pairwise synonymous substitution rate (Ks) can be used to verify the evolutionary scenarios, identify those involving hybridization with a ghost lineage, and provide a (relative) estimate of divergence time.

Target enrichment has become widely used across a diverse set of taxa in recent years, often employing the same targeted loci through universal probe sets (Johnson et al. 2019; Bartoš et al. 2023), which allows for the combination or extension of datasets in future studies. This also predisposes target enrichment phylogenomics datasets to follow-up analyses using newly developed approaches. The prevalent focus on single-copy genes often leads to filtering of phylogenomics datasets with the aim to exclude paralogous gene sequences that arose through WGDs. However, we argue that paralogs originating from such WGDs, whether through autopolyploidization or hybridization, can and should be used to gain a deeper understanding of plant evolutionary history.

Many phylogenomic studies a priori select taxa that are assumed to be diploid, or include only known and recent allopolyploids, utilizing existing bioinformatics pipelines for assembling paralogs from such taxa (Nauheimer et al. 2021; Freyman et al. 2023; Patel et al. 2023). However, the ploidy status of many plant species is still unknown (Halabi et al. 2023) and cannot easily be determined from material available only as herbarium vouchers or dried leaf material, especially older material (Viruel et al. 2019). While allele frequency-based approaches have been used to estimate ploidy levels from target enrichment data (Viruel et al. 2019), these methods may be limited to low ploidy levels up to tetraploid (Weiß et al. 2018) and were therefore not applicable to our dataset. Instead, we employed a sequence-based approach that utilizes the divergence and diversity of target enrichment loci to select diploids. Ploidy was unknown for 44% of our samples, but the high consistency between predicted and known ploidy status for the remaining samples indicated a high reliability of the method.

Unlike chromosome counts, the sequence-based approach can detect taxa with past hybridization events that have undergone diploidization, reverting their chromosome count to diploid level. However, this method may overlook recent autopolpyloids due to its reliance on allele divergence. This oversight appears to be minor in our dataset, as we identified only four samples where chromosome numbers reported in the literature suggest polyploidy, yet the respective sample was classified as ‘diploid’. Of these, three were inferred to be polyploids using kmer based ploidy inference.

Phylogenomic reconstructions of our data using conventional tree-building methods resulted in bifurcating trees largely consistent with previous single-marker phylogenetic studies (Jordon-Thaden et al. 2010; Karl and Koch 2013) which displayed high support values (posterior probability) in the backbone of Arabideae. However, they exhibit low resolution and support within clades, particularly within *Draba*, despite utilizing nearly 1,000 genes.

Specifically, we subjected our dataset to three different methods of phylogenomic tree reconstruction (Fig. 1), each of them using gene tree-species tree reconciliation but with different input data. The first tree (consensus tree) used consensus sequences of alleles or paralogs for each gene, a common approach after filtering for loci abundant with paralogs (e.g. Hendriks et al. 2023). The consensus tree showed higher support values at deeper nodes but low support within genera. This pattern likely results from low resolution in gene trees towards the tips due to ambiguity in consensus sequences, possibly caused by hybridization events, such as the one hybridization event in cluster 1 leading to the emergence of a new evolutionary lineage with over 100 taxa. The main lineages were recovered using the consensus tree, with the exception of *Arabis parvula* and *Arabis aucheri*, which previous studies showed to be sister species (Karl et al. 2012). In our analysis, *A. aucheri* grouped with *Aubrieta*, and *A. parvula* grouped with *Drabella*, possibly due to similar ploidy levels. The second tree (‘paralog tree’) included all assembled paralogs, and showed slightly lower support values than the consensus tree, but yielded phylogenetic groups consistent with the literature. The third tree (‘diploid tree’) used only seemingly diploid samples, which greatly increased support values across the tree. The major clades recovered in this subset were consistent with those in the consensus and paralog tree. Importantly, while the first two trees with all samples exhibited a caterpillar-like (pectinate) topology in most clades, in particular at nodes with low resolution within *Draba*, the samples causing this topology were largely classified as polyploids and excluded from the diploid tree.

Phylogenomic studies of clades with complex evolutionary histories often employ a multi-step approach, starting with species tree reconstruction and followed by detailed investigation of particularly interesting taxa in smaller datasets. This method is frequently justified by cytonuclear incongruence or uniform taxon sampling across clades (e.g. Morales-Briones et al. 2021; Debray et al. 2022; Fu et al. 2023). While our approach of phylogenetic placement also uses successive analyses (Fig. 1), it offers a unique advantage by allowing all data to be viewed simultaneously. This enables the assessment of all included taxa as potential origins of polyploid and hybrid taxa, providing an informative basis for subsequent selection of subsets for further analyses. For example, this approach facilitates network analyses that encompass all lineages involved in the origin of hybrid taxa, and it can also be used to inform sample selection for comparative genomics projects. Additionally, it is suitable for evolutionary scenarios that involve multiple nested WGDs, an aspect often overlooked in other phylogenomic studies.

Finally, analyzing synonymous substitution rate (*Ks*) between paralogs provided valuable insights not only into the placement of polyploid clades and the potential parents of allopolyploids, but also their relative times of origin. Within-species *Ks* is commonly used in comparative genomics studies – particularly when whole genome sequences are available – to infer the timing of WGDs across taxa (e.g. Hoang et al. 2023). It is also used in conjunction with between-species *Ks* to provide a relative timing context among related clades (Hoang et al. 2024). *Ks* has previously been employed in phylogenetic studies to assess the number of WGDs and compare patterns of (shared) *Ks* peak distribution based on whole-genome and transcriptome data (Morales-Briones et al. 2021). In our study, we calculated pairwise *Ks* between samples and within polyploids to evaluate the presence of past WGDs, as well as patterns and order of divergence. We were able to assess *Ks* within and between all samples from target enrichment data directly and successfully detected and distinguished three modes of polyploidization: Hybridization (allopolyploidization), autopolyploidization, as well as hidden allopolyploidization involving a ghost lineage. The latter, in particular, only became apparent through the interpretation of within- and between-species *Ks* peaks, and is generally difficult to discern using phylogenetic methods. Thus, ghost lineages are largely overlooked as they cannot be detected using traditional phylogenetic tree reconstruction methods. Unlike introgression, ghost lineages may have only a small impact on divergence time estimates unless a significant number of loci were retained from the respective hybridization event (Tiley et al. 2023). Recently, methods derived from population genetics, such as coalescent modeling, or comparative genomics have been employed to hypothesize or detect ghost lineages in specific scenarios (Luo et al. 2017; Ru et al. 2018; Dong et al. 2022; Sancho et al. 2022; Chang et al. 2023). Network analyses have also been successful in reconstructing evolutionary scenarios involving ghost lineages as the parents of even multiple hybrid samples in *Erysimum* (Osuna-Mascaró et al. 2023). These studies altogether underscore the significant role that ghost lineages can play in the evolutionary history of a various taxa and highlight the challenges associated with their detection. The use of *Ks* for discovering such ghost lineages adds another layer of evidence to support such events and could be a valuable tool in other studies.

In summation, there is no one-size-fits-all approach to phylogenomics of hybrids and polyploids; different datasets may require different strategies. For instance, when genome sequences are available, subgenomes analysis may be a viable option (Guo et al. 2021). Our method of Paralog PhyloGenomics offers first insights into the evolutionary processes that have shaped phylogenetically complex and highly polyploid clades using target enrichment data (and could easily be extended to other multi-locus datasets). These initial insights can and should serve as a foundation for further genome-scale studies to develop a more comprehensive understanding of the intricate patterns of reticulate evolution found in plants. Moreover, our results highlight the necessity for developing methods to study trait and character evolution in the context of hybridization and polyploidization (Wang et al. 2021). Bifurcating trees fail to capture the full evolutionary reality of such clades, and their evolutionary potential may be influenced by contributions from multiple lineages.

Oversimplification into phylogenetic trees can significantly limit the conclusions drawn from comparative phylogenetic or biogeographic studies. Therefore, it is essential to adopt methodologies that account for the complexity and reticulate nature of plant evolution to fully understand their evolutionary history.

## Supplementary Tables and Figures

**Table S1. Sample and sequencing information. Table S2. HybPhaser gene and sample recovery.**

**Figure S1. Mapping rate.**

**Figure S2. Gene recovery rate.**

**Figure S3. Gene selection.**

**Figure S4. Locus heterozygosity and allele divergence.**

**Figure S5. Allele divergence (AD) and locus heterozygosity (LH).**

**Figure S6. Comparison of different ploidy inference methods.**

**Figure S7. Number of paralogs.**

**Figure S8. Paralog detection by ploidy.**

**Figure S9. ASTRAL tree based on consensus sequences.**

**Figure S10. Support values for ASTRAL tree based on consensus sequences.**

**Figure S11. ASTRAL-pro tree based on paralog sequences.**

**Figure S12. Support values for ASTRAL-pro tree based on paralog sequences.**

**Figure S13. ASTRAL-pro tree based on paralogs from diploid samples.**

**Figure S14. Support values for ASTRAL-pro tree based on paralogs from diploid samples.**

**Figure S15. Gene selection for phylogenetic placement.**

**Figure S16. Phylogenetic placement.**

**Figure S17. Scatterplot of clustering height and number of clusters.**

**Figure S18. Dendrogram of hierarchical clustering.**

**Figure S19. Dendrograms for paralog placement of clusters 1-4.**

**Figure S20. Dendrograms for paralog placement of clusters 5-8.**

**Figure S21. Dendrograms for paralog placement of clusters 9-12.**

**Figure S22. Dendrograms for paralog placement of clusters 13-16.**

**Figure S23. Dendrograms for paralog placement of clusters 17-20.**

**Figure S24. Dendrograms for paralog placement of clusters 21-24.**

**Figure S25. Dendrograms for paralog placement of clusters 25-28.**

**Figure S26. Dendrograms for paralog placement of clusters 29-32.**

**Figure S27. Dendrograms for paralog placement of clusters 33-36.**

**Figure S28. Dendrograms for paralog placement of clusters 37-40.**

**Figure S29. Dendrograms for paralog placement of clusters 41-43.**

**Figure S30. Three evolutionary scenarios for the emergence of polyploids analyzed with Ks.**

**Figure S31. Synonymous substitution rate distribution in a clade with hybridization.**

**Figure S32. Synonymous substitution rate distribution in a clade with WGD.**

**Figure S33. Synonymous substitution rate distribution in a clade with WGD.**

**Figure S34. Synonymous substitution rate distribution in a clade with a ghost lineage.**

## Acknowledgements

We are grateful to Ihsan Al-Shehbaz and Dmitry German for their continuous and generous support and many helpful comments on taxonomy, Markus Kiefer for bioinformatics assistance, Anna Loreth for DNA extraction and library preparation, and Peter Sack for assistance with herbarium vouchers and collection management. We also thank Kasper Hendriks, Klaus Mummenhoff and Frederic Lens for granting early access to data from Hendriks et al. (2023), and Eric Schranz and Dan Kliebenstein for helpful comments on the draft version.

## Data and Supplementary Materials

Raw sequencing reads generated for this study are available at ENA (https://www.ebi.ac.uk/ena/browser/home) under accession PRJEB72669.

