## Supplementary Material for "Unravelling complex hybrid and polyploid evolutionary relationships using phylogenetic placement of paralogs from target enrichment data"

*Corresponding authors:*

### Supplementary Tables

#### Table S2. HybPhaser gene and sample recovery.

Our selection of loci and samples was compared to filtering strategies 'inclusive', 'strict' and 'superstrict' from (Hendriks et al. 2023).

| | All selected loci and samples | Inclusive | Strict | Superstrict |
|---|---|---|---|---|
| **Gene and sample recovery thresholds** | | | | |
| Gene minimum proportion of samples recovered | 0 | 0.1 | 0.2 | 0.2 |
| Gene minimum proportion of target length recovered | 0 | 0.1 | 0.2 | 0.2 |
| Sample minimum proportion of total target length recovered | 0 | 0 | 0.4 | 0.4 |
| Sample minimum proportion of genes recovered | 0 | 0 | 0.2 | 0.2 |
| Gene SNPs proportion threshold for all samples | none | none | outliers | 0.02 |
| Remove outlier genes per sample? | no | yes | yes | yes |
| **Results** | | | | |
| Number of genes retained | 994 | 978 | 912 | 34 |
| Number of samples | 442 | 438 | 438 | 438 |

### Supplementary Figures

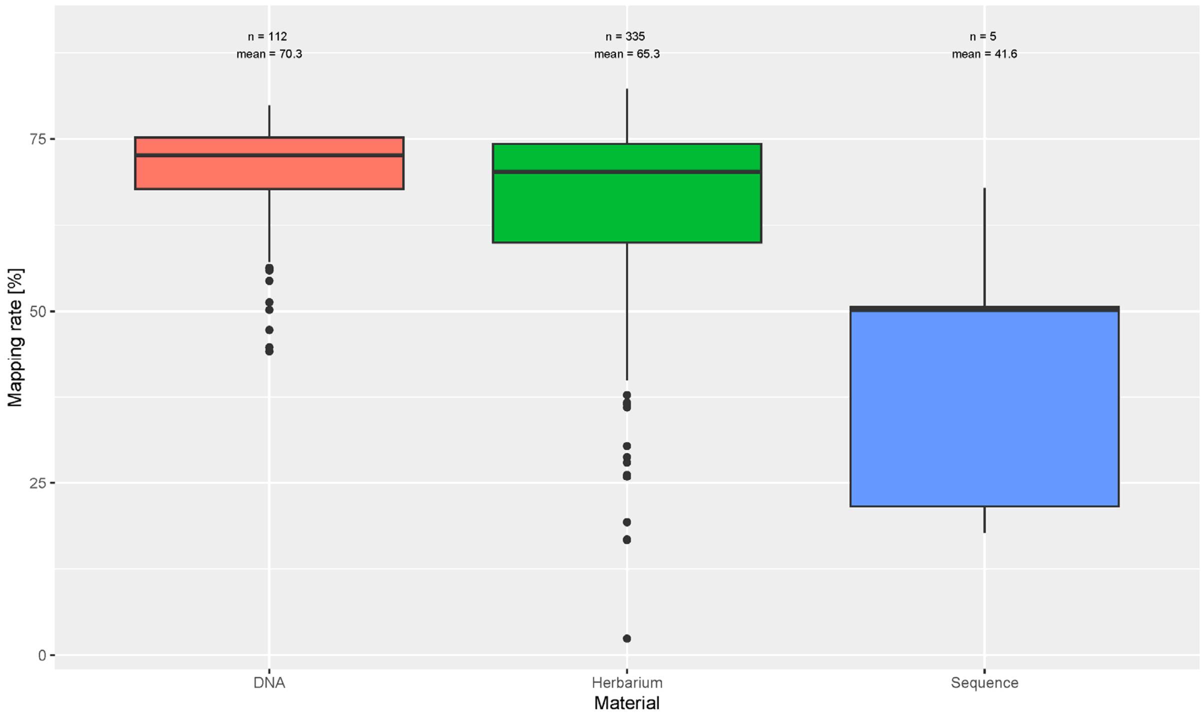

**Figure S1. Mapping rate.** Percentage of reads mapped to the target reference is given for each source material. New sequences were generated for this study either from DNA used in previous studies or from DNA extracted from herbarium vouchers. Additionally, sequences from five samples were added from Hendriks et al. (2023).

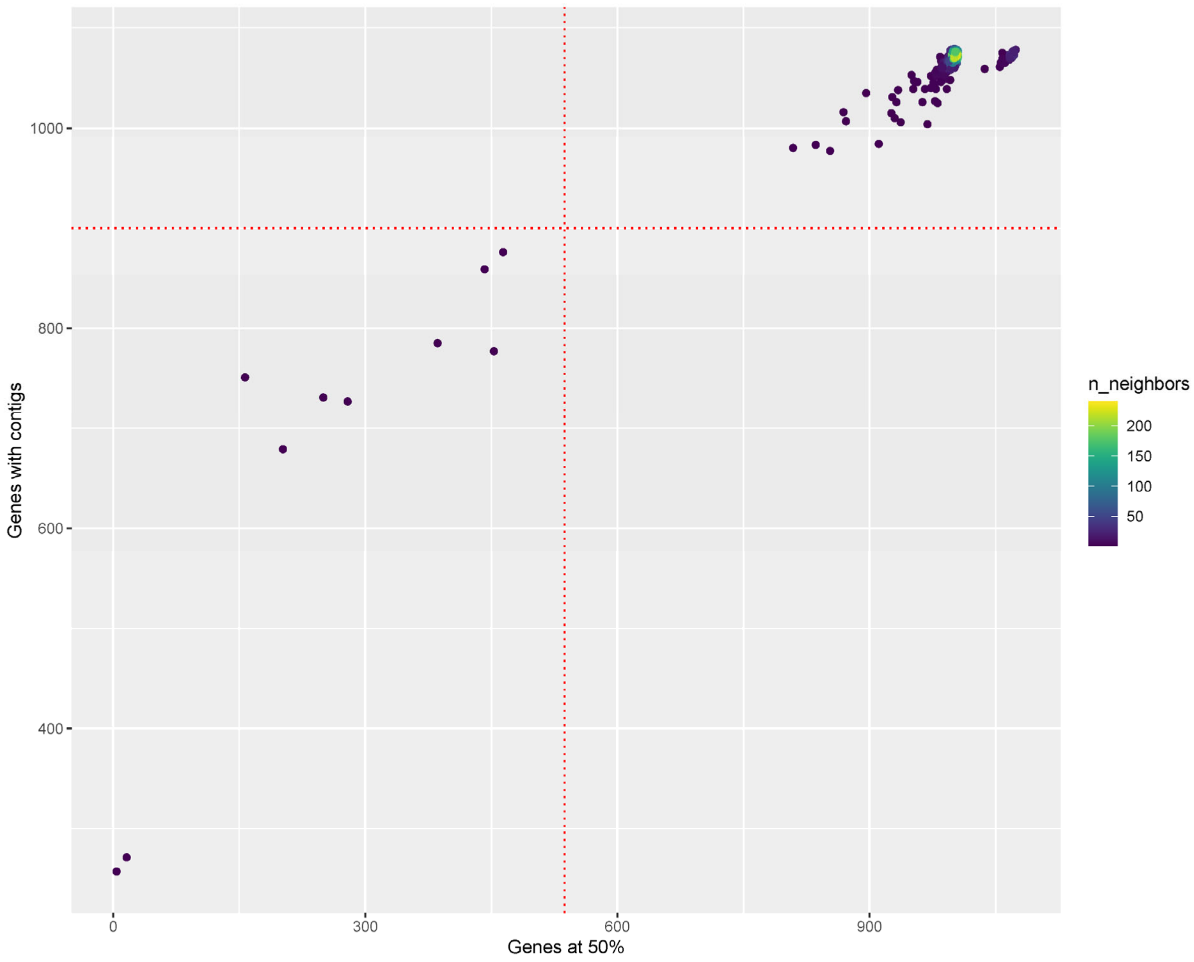

**Figure S2. Gene recovery rate.** Point density plot for the number of genes with contigs and number of genes covered over a minimum length of 50% of the target reference. Only samples with at least 900 genes with contigs and half of the total genes covered at 50% were retained for further analysis; the selection threshold is indicated by red dotted lines. The final dataset thus comprised 442 samples.

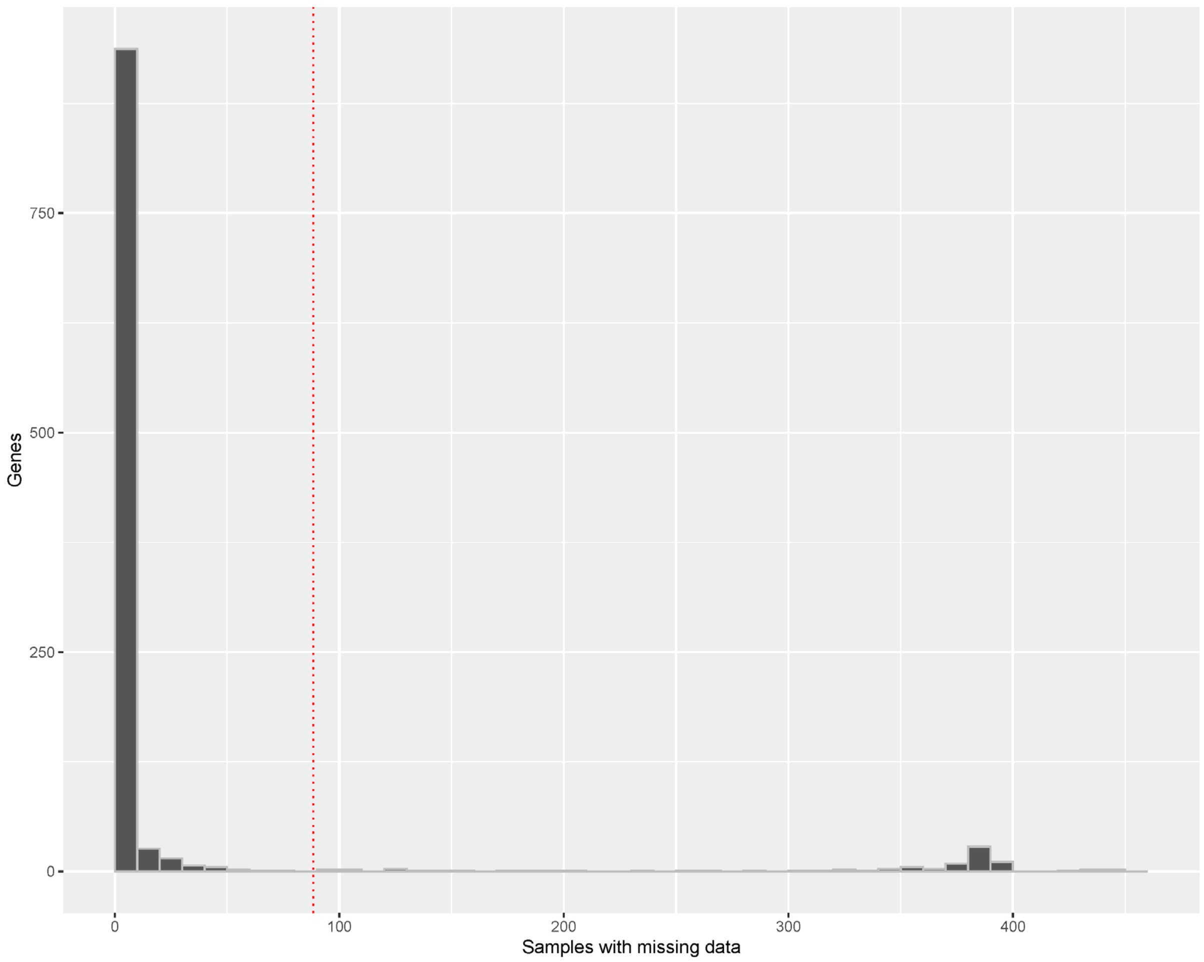

**Figure S3. Gene selection.** Histogram of number of genes and samples with missing data. Only genes with less than 20% of samples missing were retained for further analysis; the selection threshold is indicated by red dotted lines. The final dataset thus comprised 994 genes.

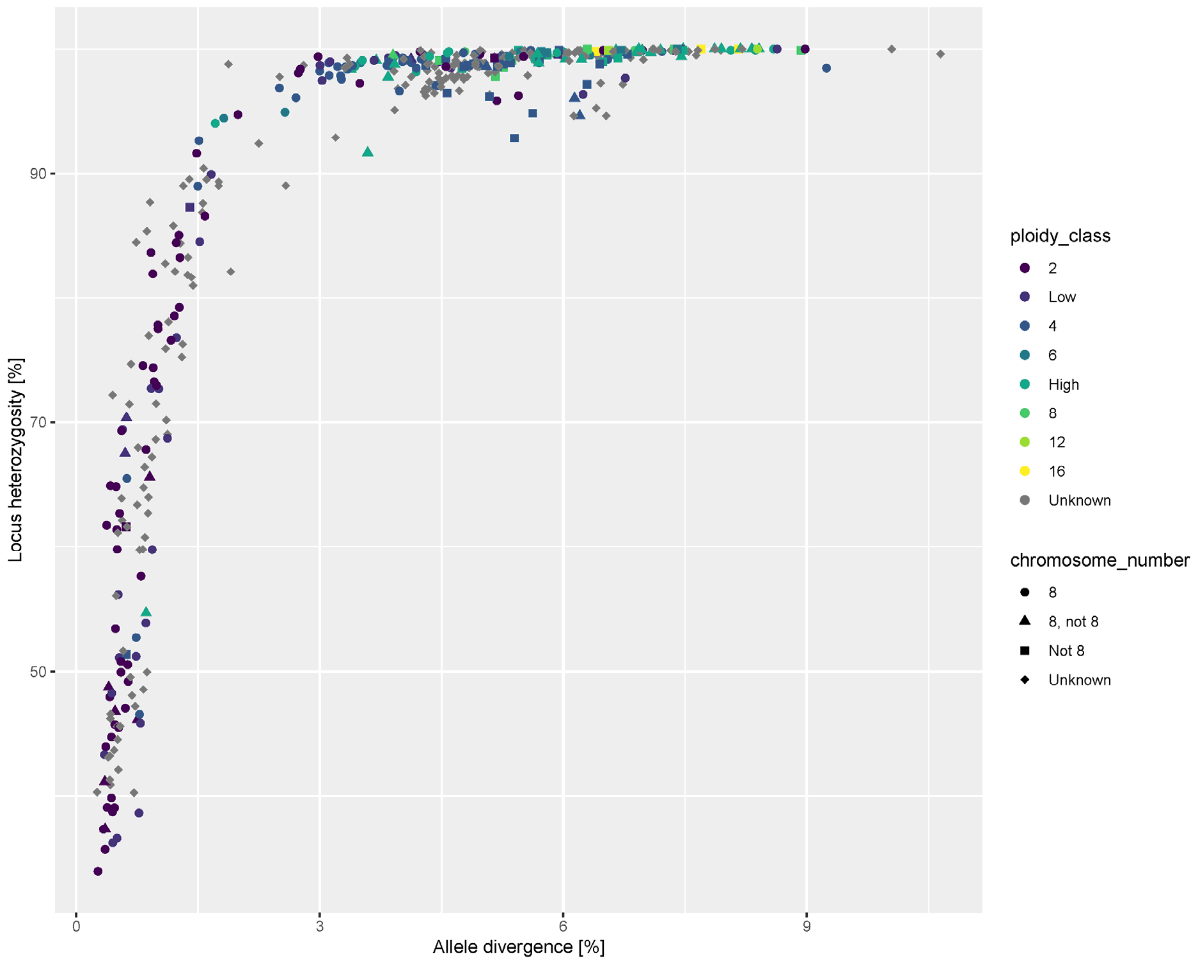

**Figure S4. Locus heterozygosity and allele divergence.** HybPhaser results for locus heterozygosity (LH) and allele divergence (AD) are shown with published ploidy level and chromosome number. Taxa with high ploidy levels generally showed high locus heterozygosity and allele divergence, in line with expectations.

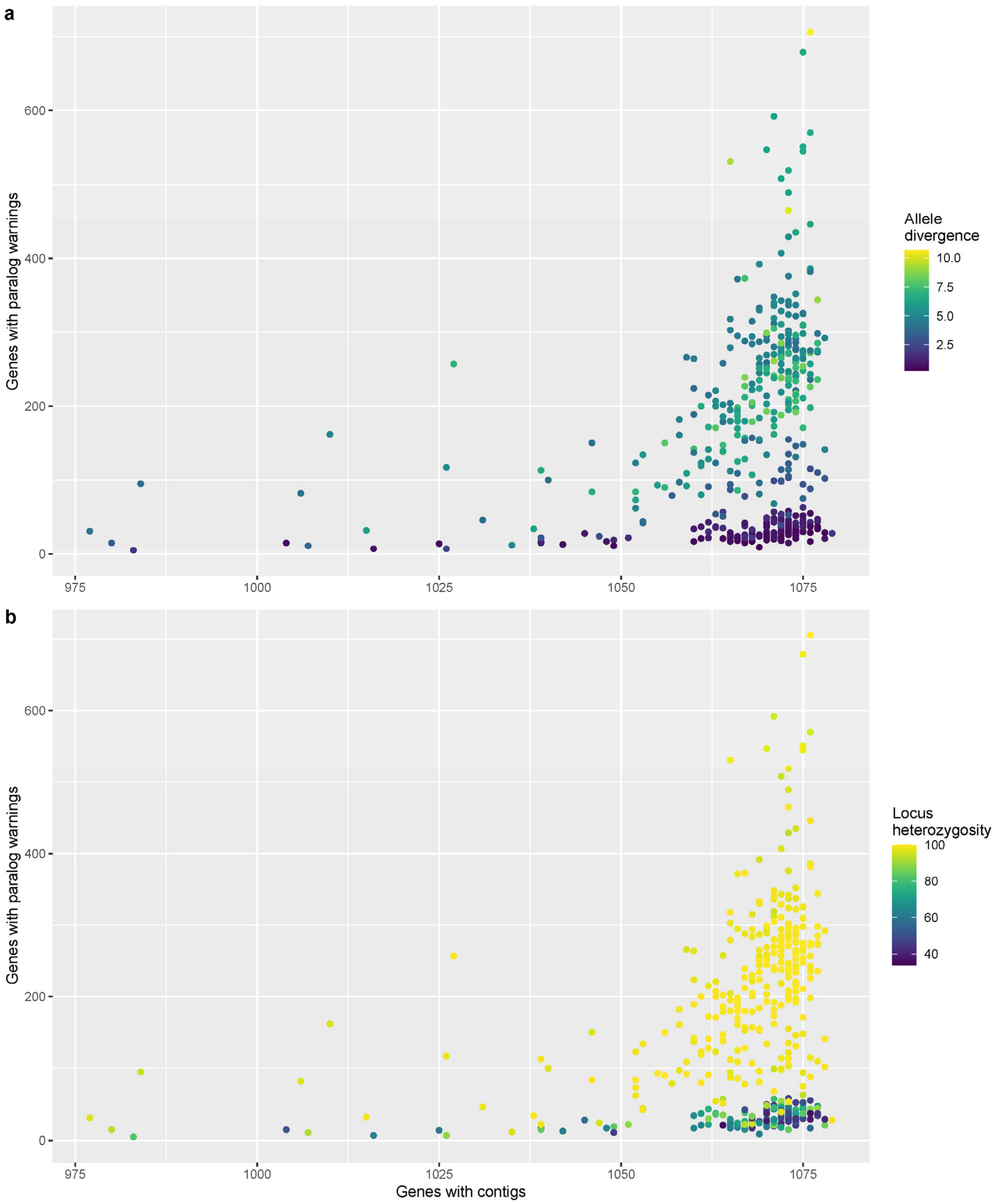

**Figure S5. Allele divergence (AD) and locus heterozygosity (LH).** (a) AD of the different paralogs within each sample as obtained by HybPhaser is plotted against the number of paralog warnings from HybPiper and the number of contigs with sequences. (b) LH of the different paralogs within each sample as obtained by HybPhaser is plotted against the number of paralog warnings from HybPiper and the number of contigs with sequences. Samples that failed our minimum sequence recovery threshold of 900 genes and genes that failed our minimum sequence recovery threshold of less than 20% missing samples were excluded from the analysis. AD and LH were used to select 138 'diploid' individuals for separate phylogenomic reconstruction (AD < 2 and LH < 90).

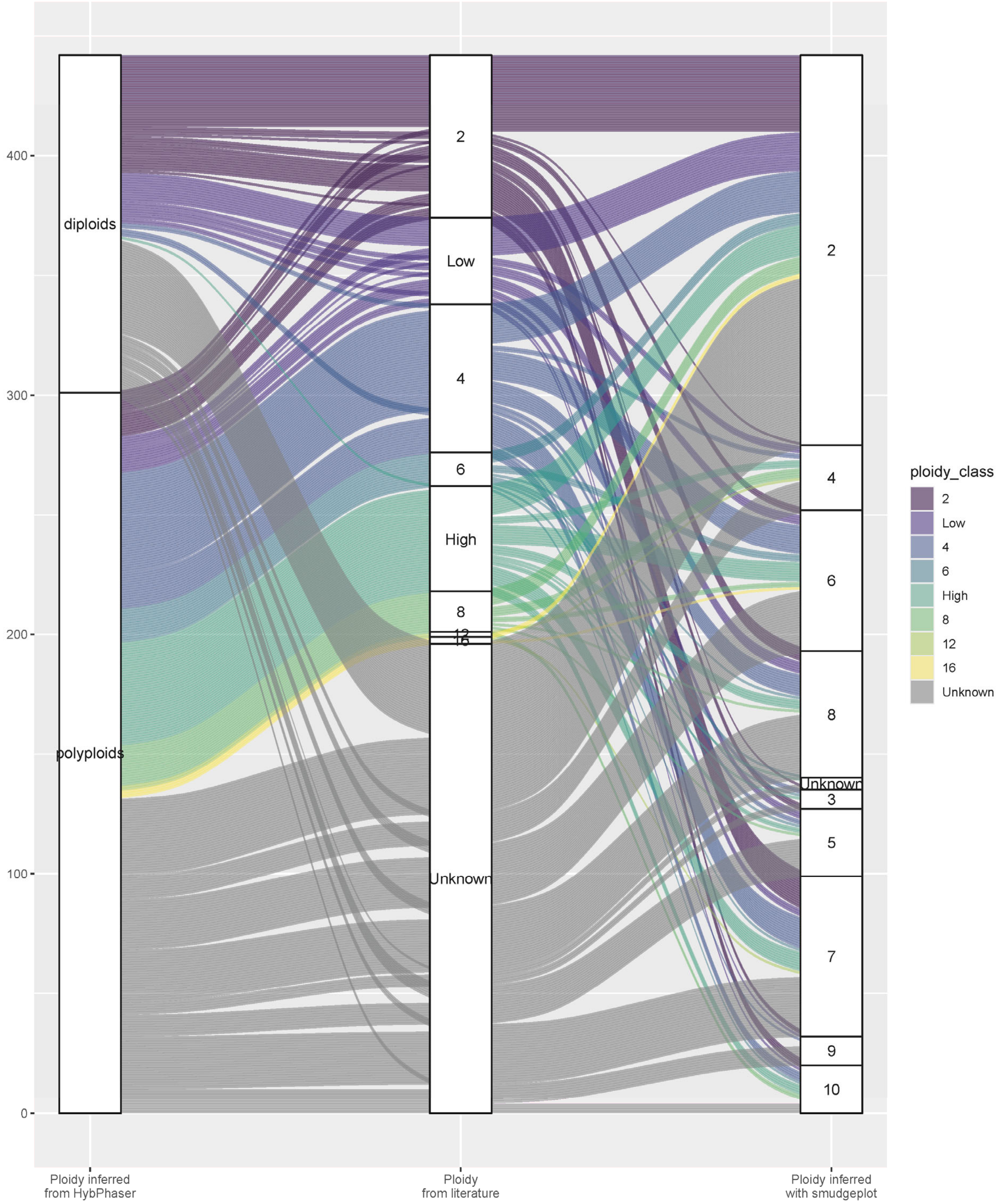

**Figure S6. Comparison of different ploidy inference methods.** Alluvial plot illustrating the results of ploidy inference using HybPhaser and smudgeplot in relation to ploidy reported in the literature. Samples assigned to the 'diploids' group using HybPhaser contained mostly diploid taxa or those with low ploidy levels according to literature, while most tetraploid and hexaploid taxa and all samples with known high ploidy (> hexaploid) were classified as 'polyploids'. Conversely, ploidy assignment using smudgeplot was largely inconsistent with ploidy reported in literature, though interestingly many known diploids assigned to 'polyploids' using HybPhaser were also inferred to be polyploid.

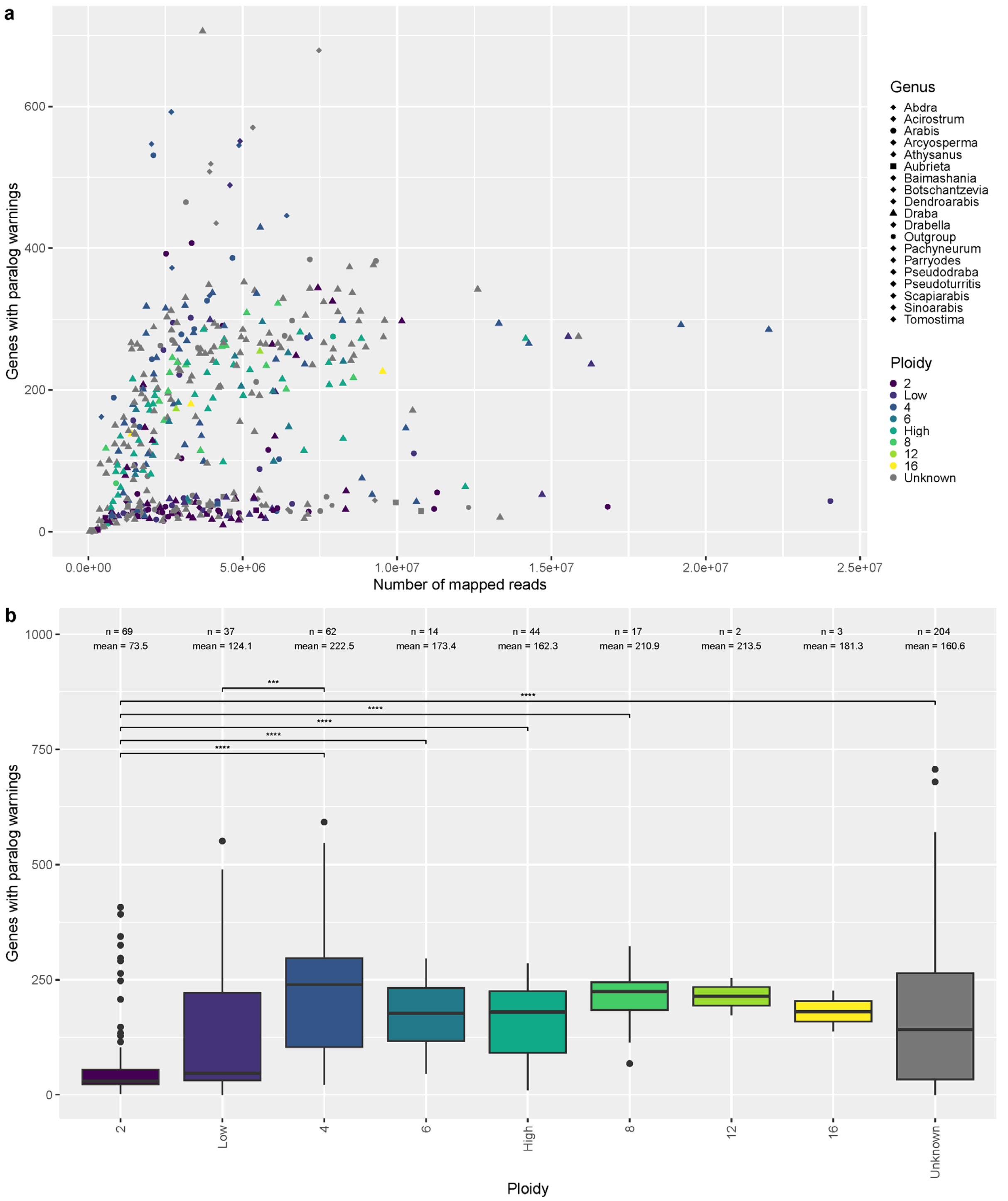

**Figure S7. Number of paralogs.** (a) Scatterplot of number of genes with paralog warnings and number of mapped reads. The larger genera are indicated with symbols, ploidy levels as reported in the literature using the color scale. (b) Boxplot of number of genes with paralog warnings by ploidy level reported in the literature. A Kruskal-Wallis test revealed significant differences between groups (Kruskal-Wallis chi-squared = 62.163, df = 8, p-value = 1.751e-10); only significant pairwise Wilcoxon rank-sum test are shown (***: p <= 0.001; ****: p <= 0.0001).

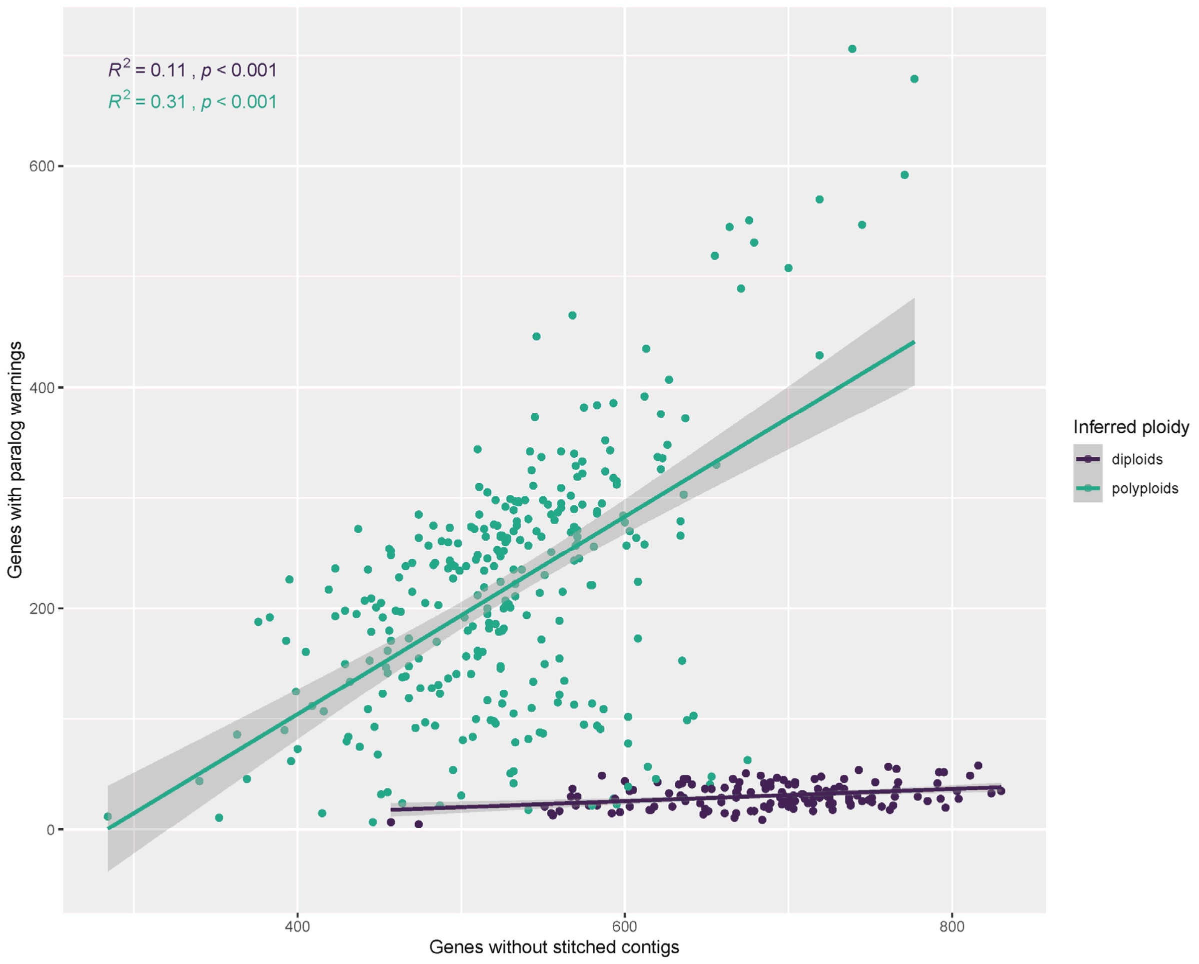

**Figure S8. Paralog detection by ploidy.** Scatterplot of the number of genes with paralog warnings from HybPiper and the number of genes without stitched contigs (i.e. those where gene sequences could be assembled as a single contig in SPAdes).

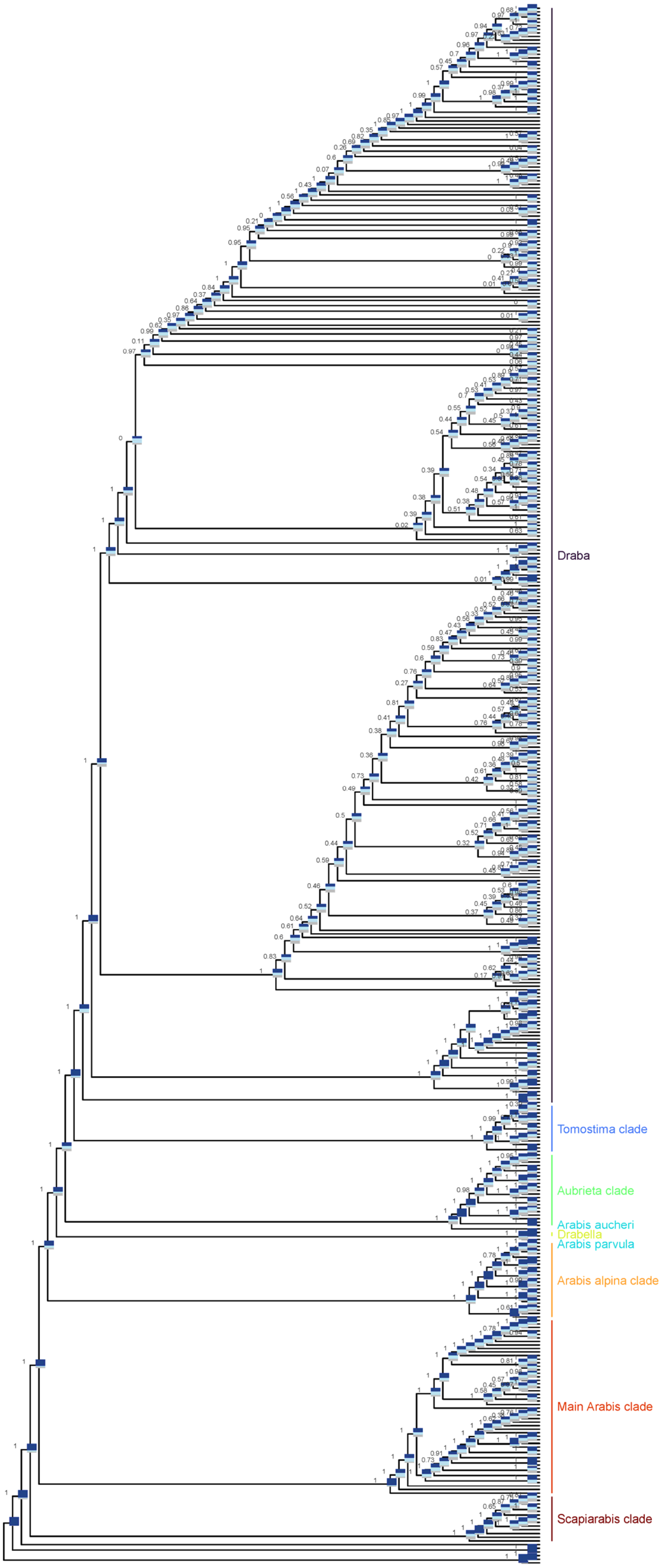

**Figure S9. ASTRAL tree based on consensus sequences.** The tree was reconstructed from 994 gene trees based on consensus sequences obtained from HybPhaser. Posterior probabilities for the first topology are shown as node labels and quartet scores are shown as bar charts: dark blue for the first quartet, light blue for the first alternative quartet, and grey for the second alternative quartet. Major Arabideae clades are annotated on the right.

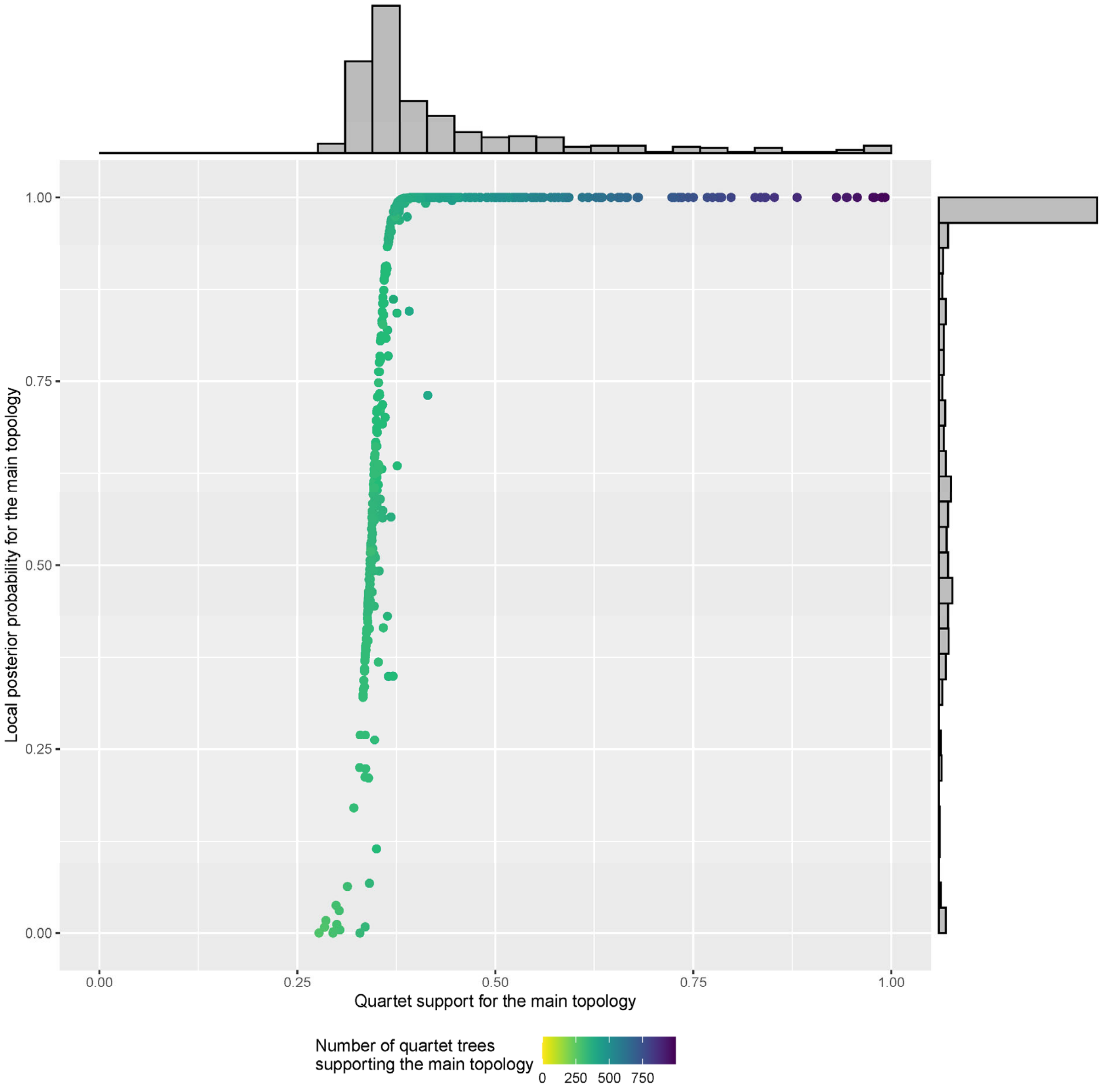

**Figure S10. Support values for ASTRAL tree based on consensus sequences.** Posterior probability and quartet support at each branch are shown for the first topology; color scale shows the number of quartet trees in all the gene trees that support the first topology relative to the total number of genes (994).

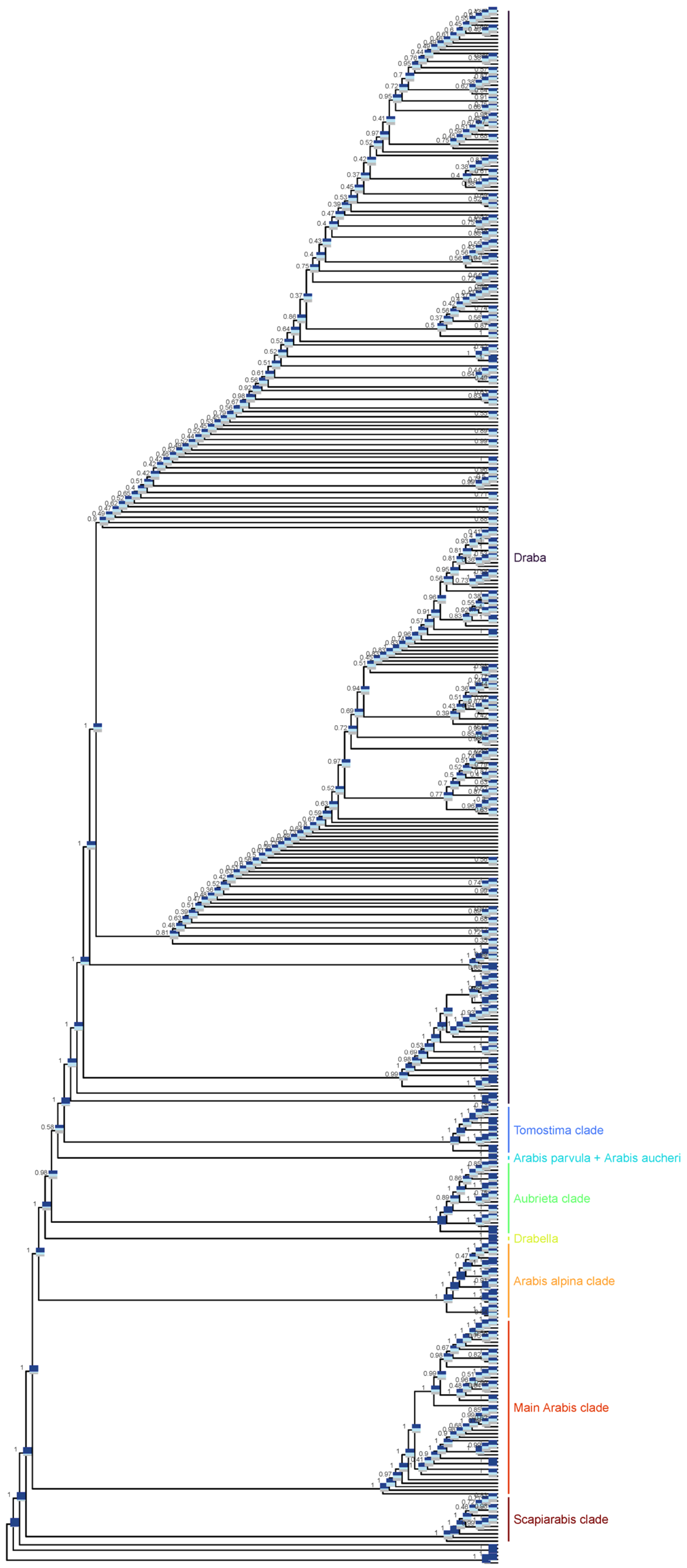

**Figure S11. ASTRAL-pro tree based on paralog sequences.** The tree was reconstructed from 994 gene trees that included all paralogs assembled by HybPiper. Posterior probabilities for the first topology are shown as node labels and quartet scores are shown as bar charts: dark blue for the first quartet, light blue for the first alternative quartet, and grey for the second alternative quartet. Major Arabideae clades are annotated on the right.

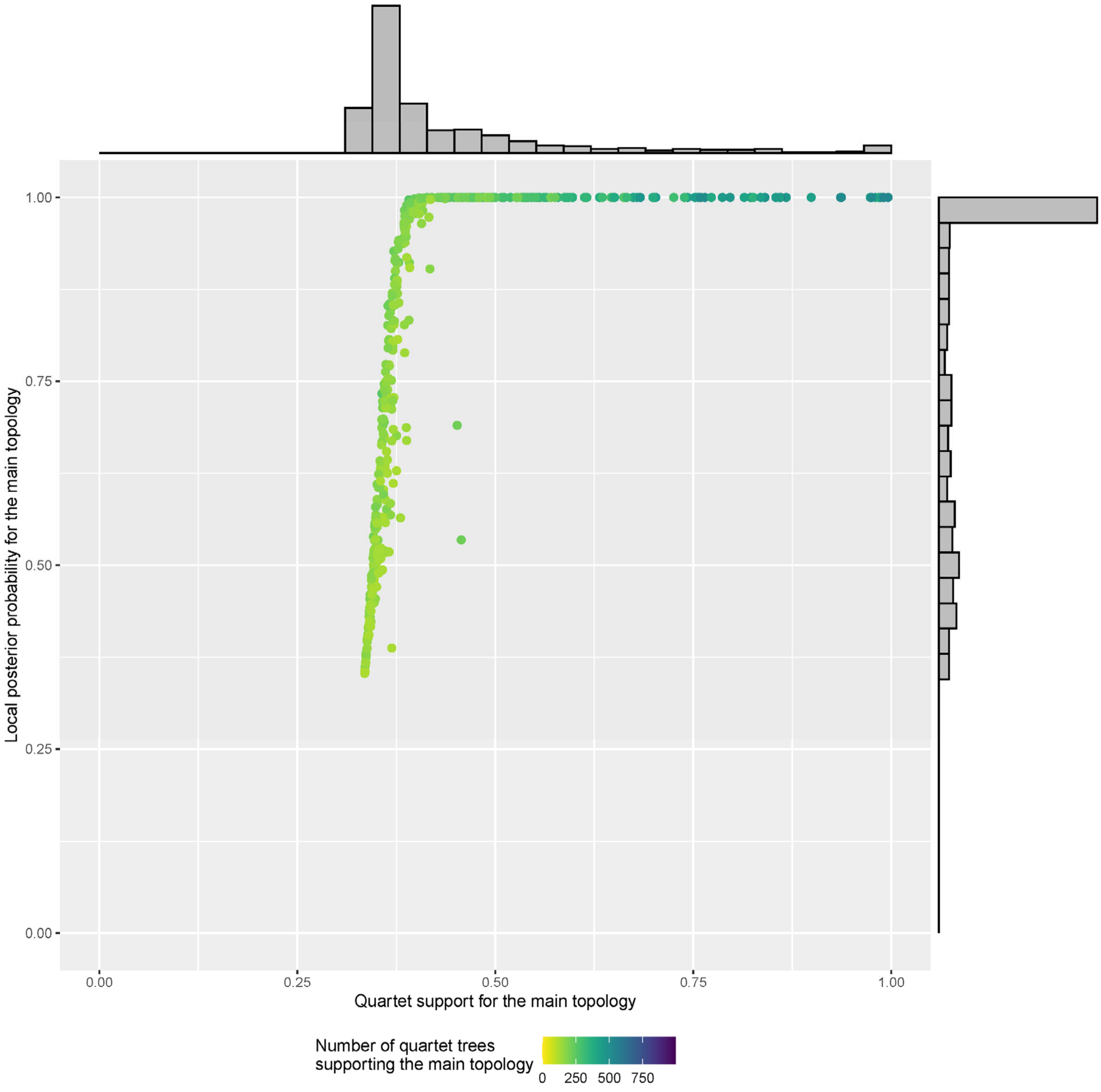

**Figure S12. Support values for ASTRAL-pro tree based on paralog sequences.** Posterior probability and quartet support at each branch are shown for the first topology; color scale shows the number of quartet trees in all the gene trees that support the first topology relative to the total number of genes (994).

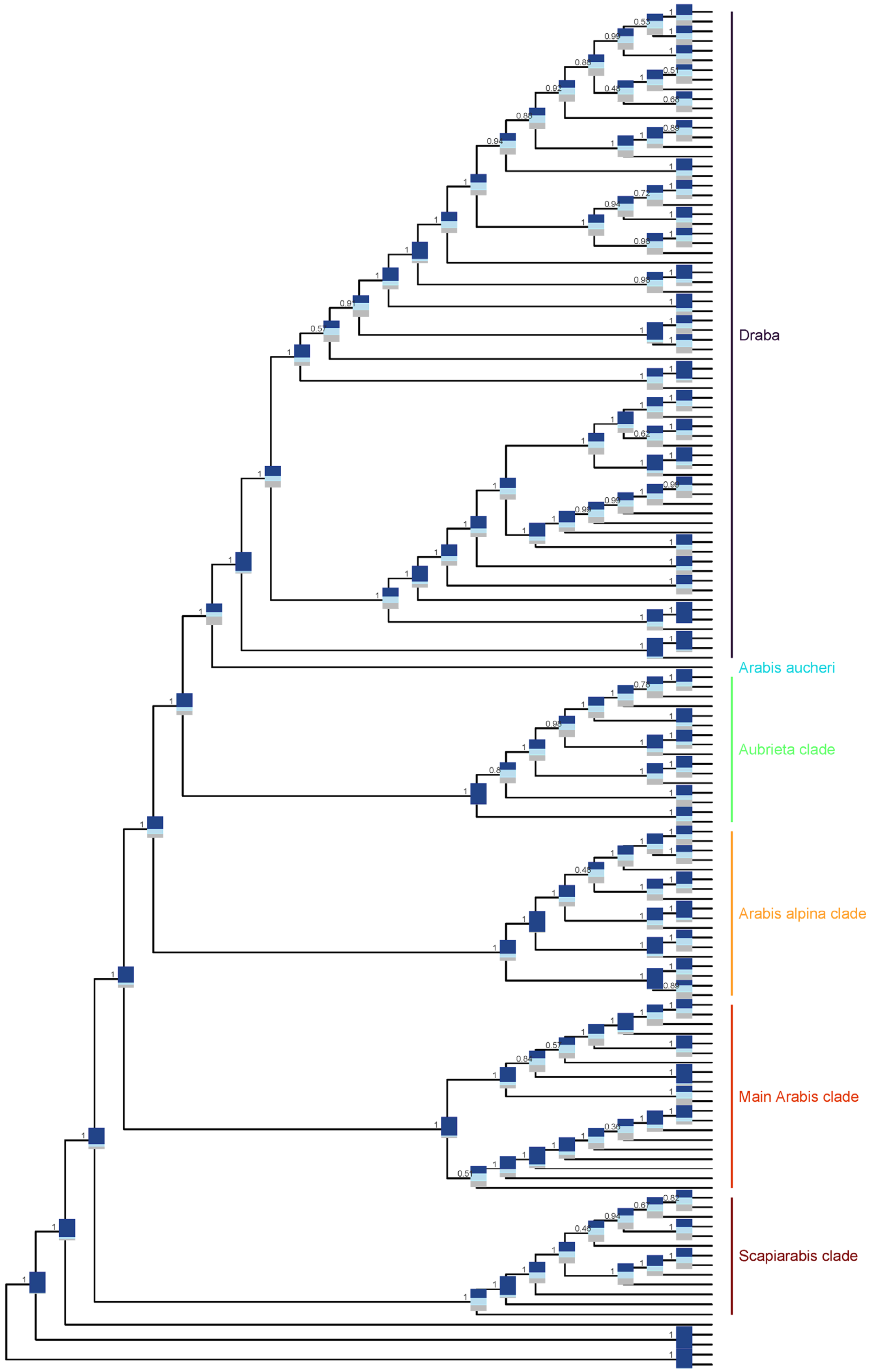

**Figure S13. ASTRAL-pro tree based on paralogs from diploid samples.** The tree was reconstructed from 994 gene trees that included all paralogs assembled by HybPiper. Posterior probabilities for the first topology are shown as node labels and quartet scores are shown as bar charts: dark blue for the first quartet, light blue for the first alternative.

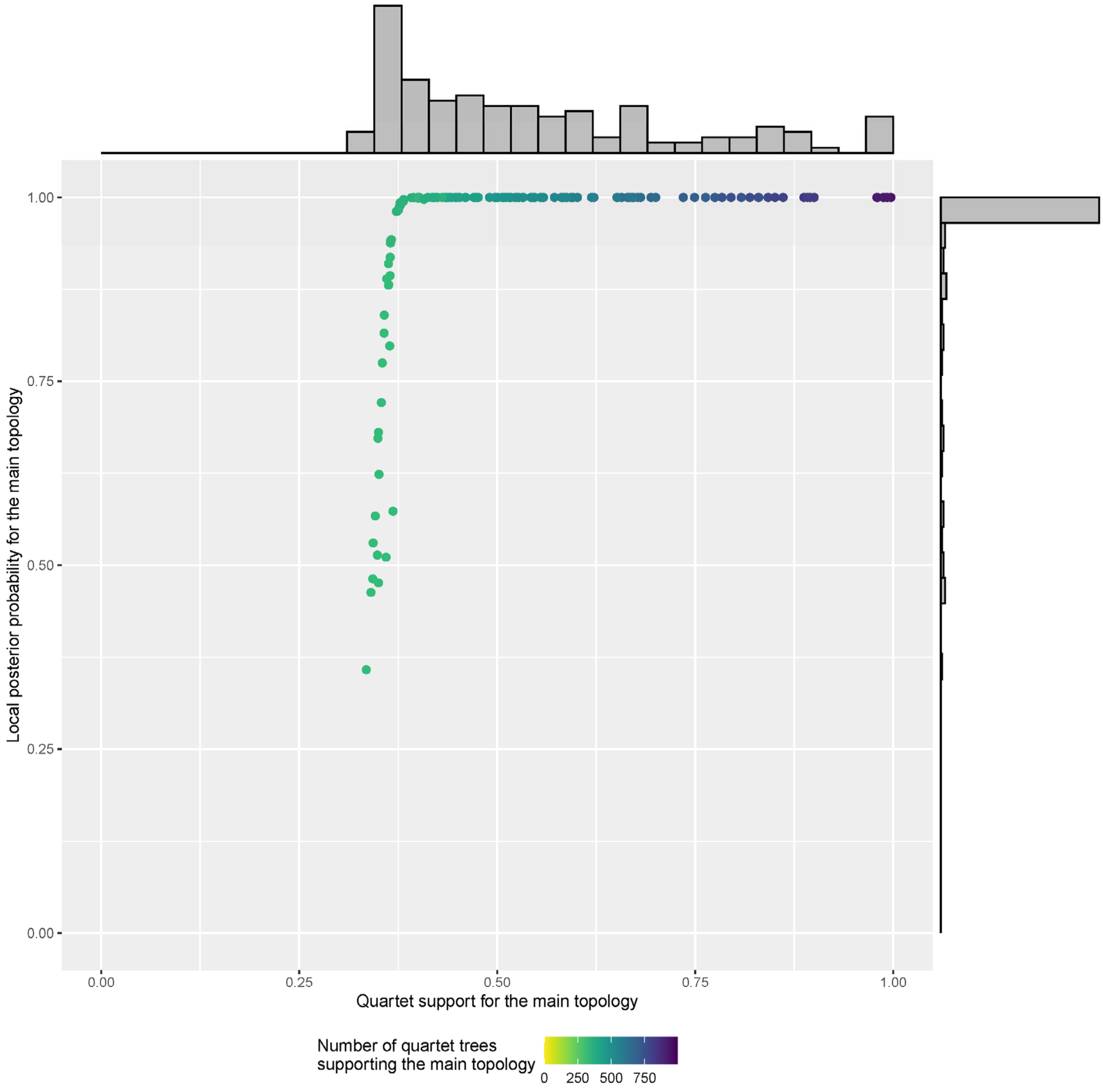

**Figure S14. Support values for ASTRAL-pro tree based on paralogs from diploid samples.** Posterior probability and quartet support at each branch are shown for the first topology; color scale shows the number of quartet trees in all the gene trees that support the first topology relative to the total number of genes (994).

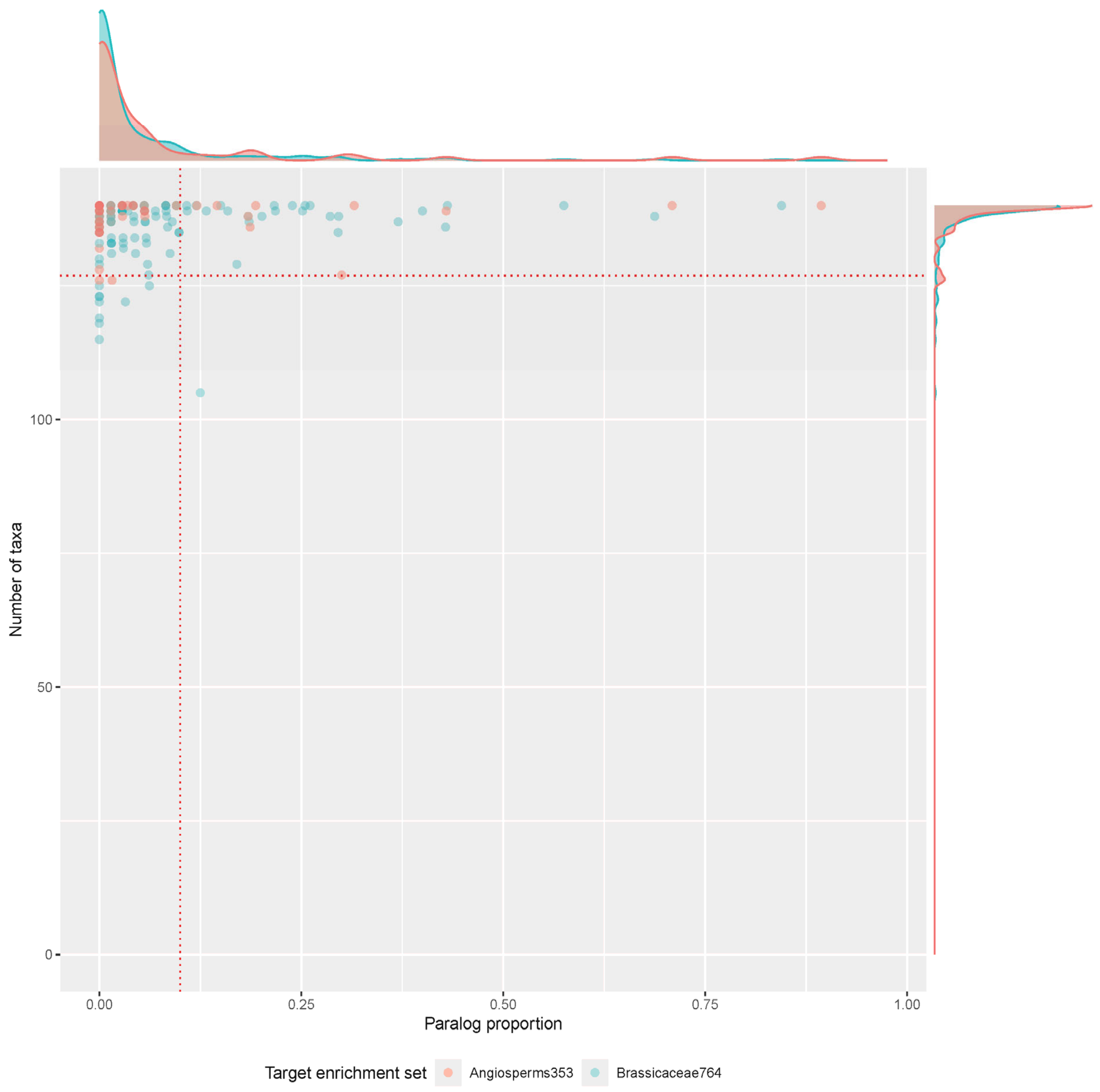

**Figure S15. Gene selection for phylogenetic placement.** Proportion of paralogs and number of taxa (of 138) is shown. Only genes with less than 10% of samples with paralogs and at least 90% of samples present were used.

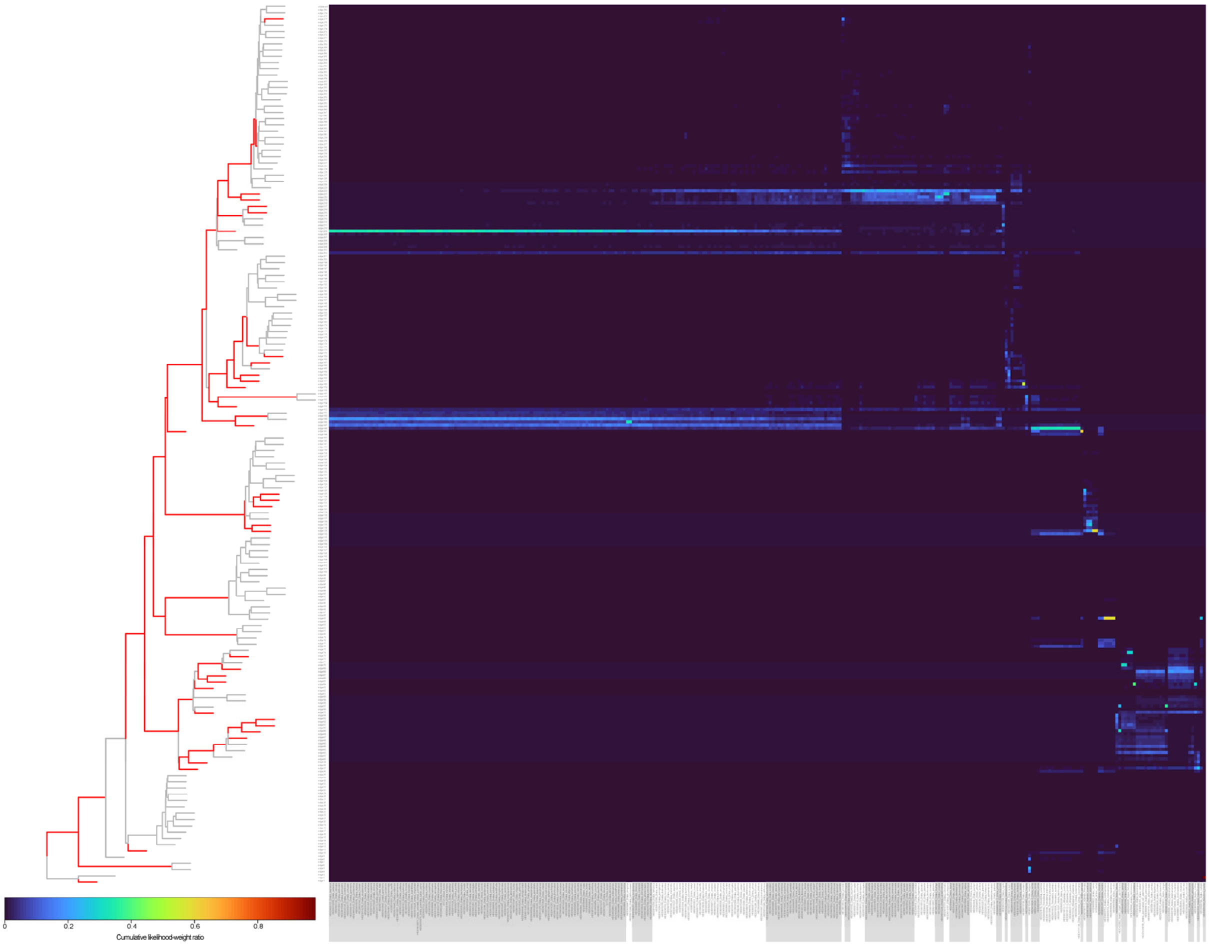

**Figure S16. Phylogenetic placement.** Cumulative likelihood-weight ratios (lwr) over all gene copies from ‘polyploid’ samples are given. Samples are arranged by cluster on the x-axis, and edges in the tree are displayed on the y-axis. The dendrogram of the diploid tree was added for orientation; edges with mean lwr ≥ 0.05 in at least one cluster are highlighted in red.

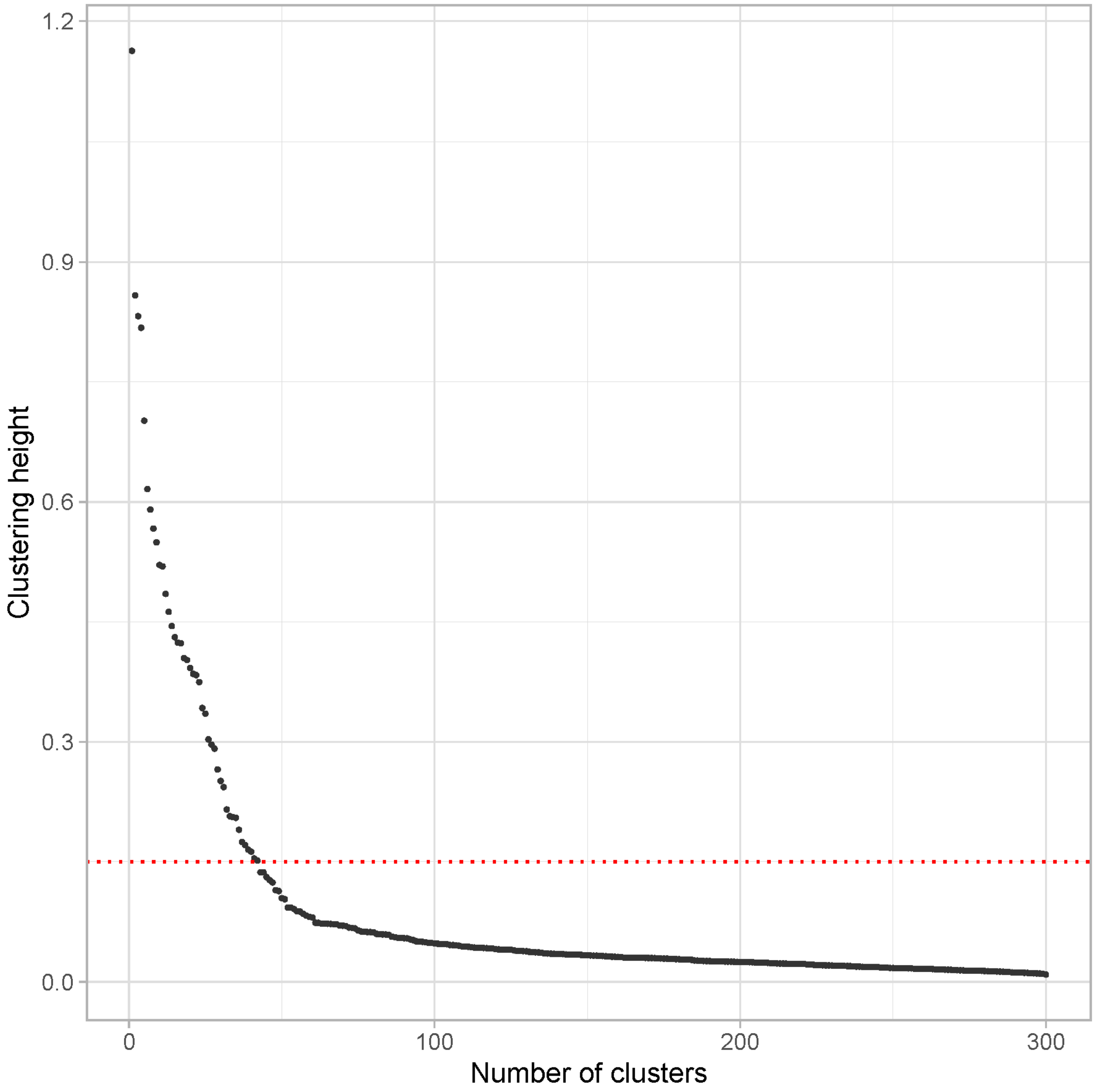

**Figure S17. Scatterplot of clustering height and number of clusters.** Distribution of clustering heights after hierarchical clustering of distance between normalized paralog placement assignment. A threshold of 0.15 was chosen to select clusters and is indicated by the red dashed line.

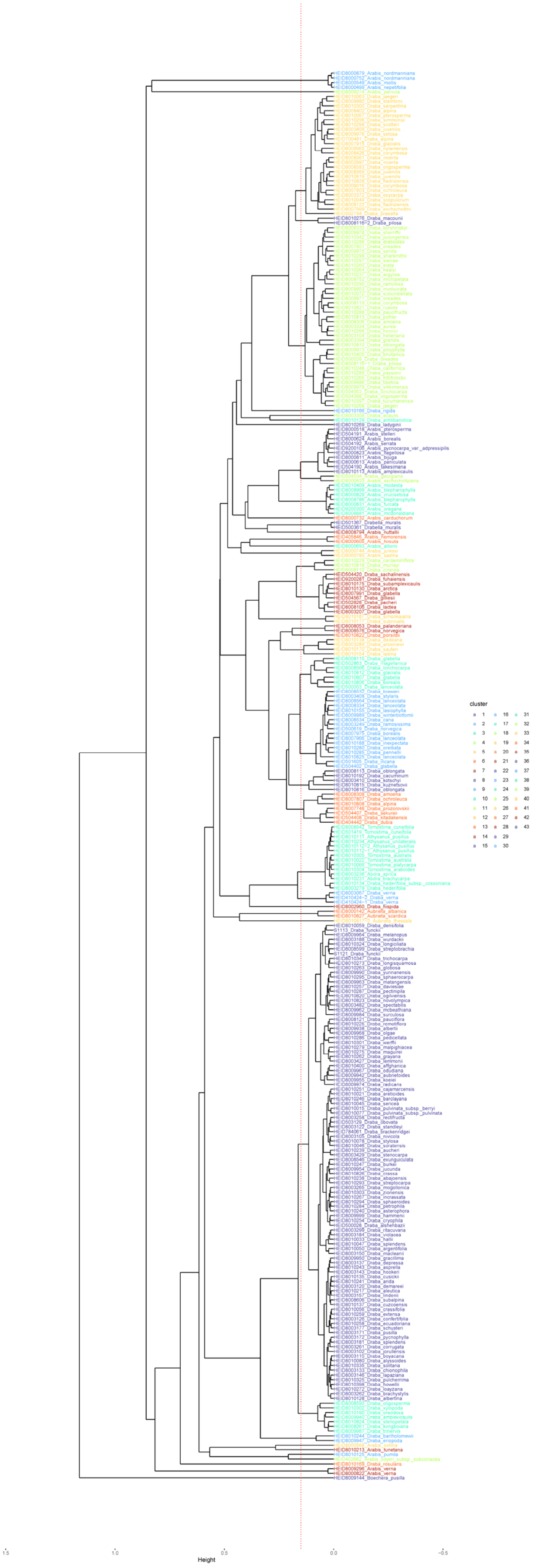

**Figure S18. Dendrogram of hierarchical clustering.** The threshold for selecting the number of clusters (0.15) is indicated by the red dashed line.

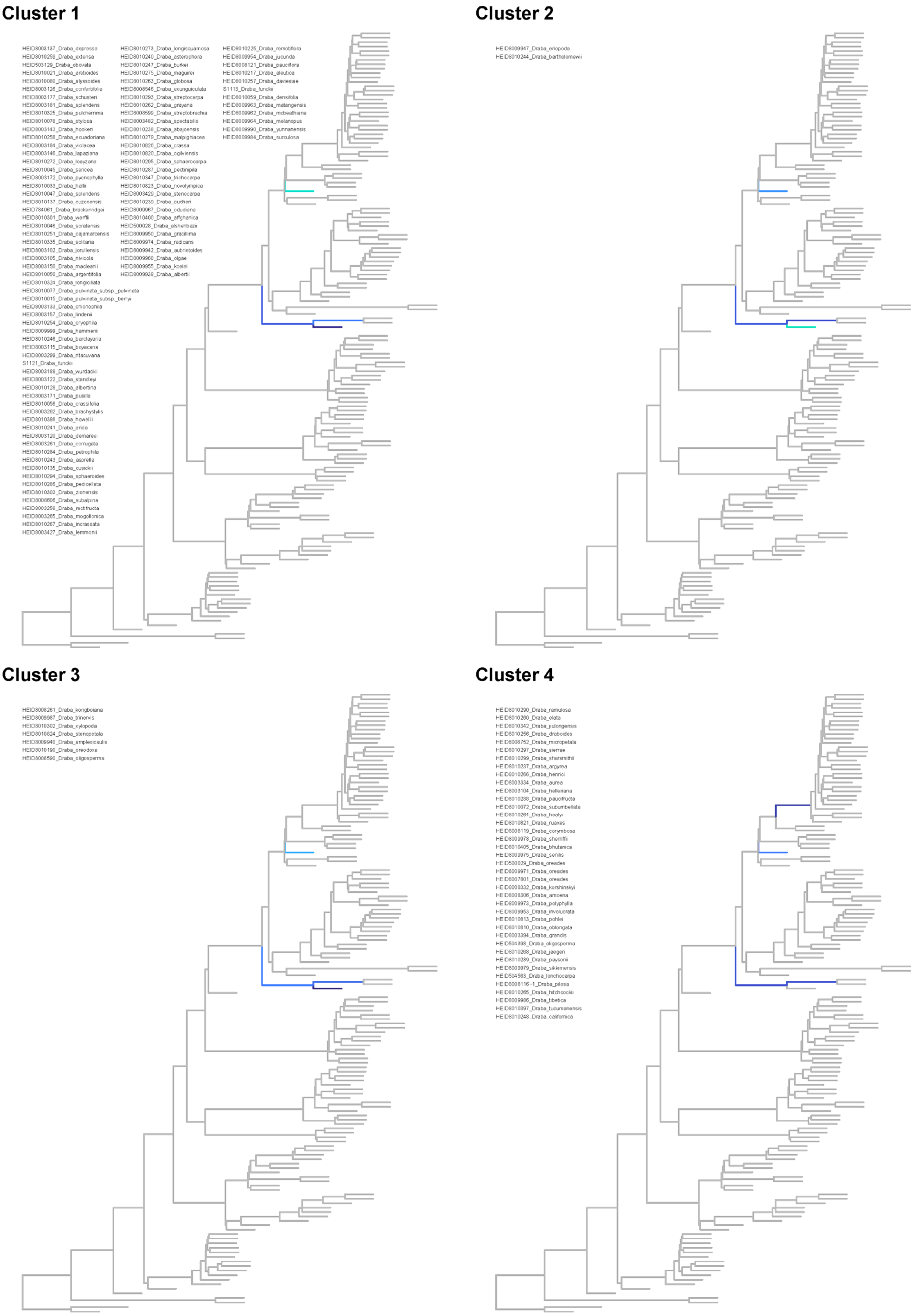

**Figure S19. Dendrograms for paralog placement of clusters 1-4.** Diploid trees with branches colored according to placement proportion are shown; samples assigned to each cluster are given in the top left corner of each subplot. A threshold of 0.05 was chosen to display placement proportion.

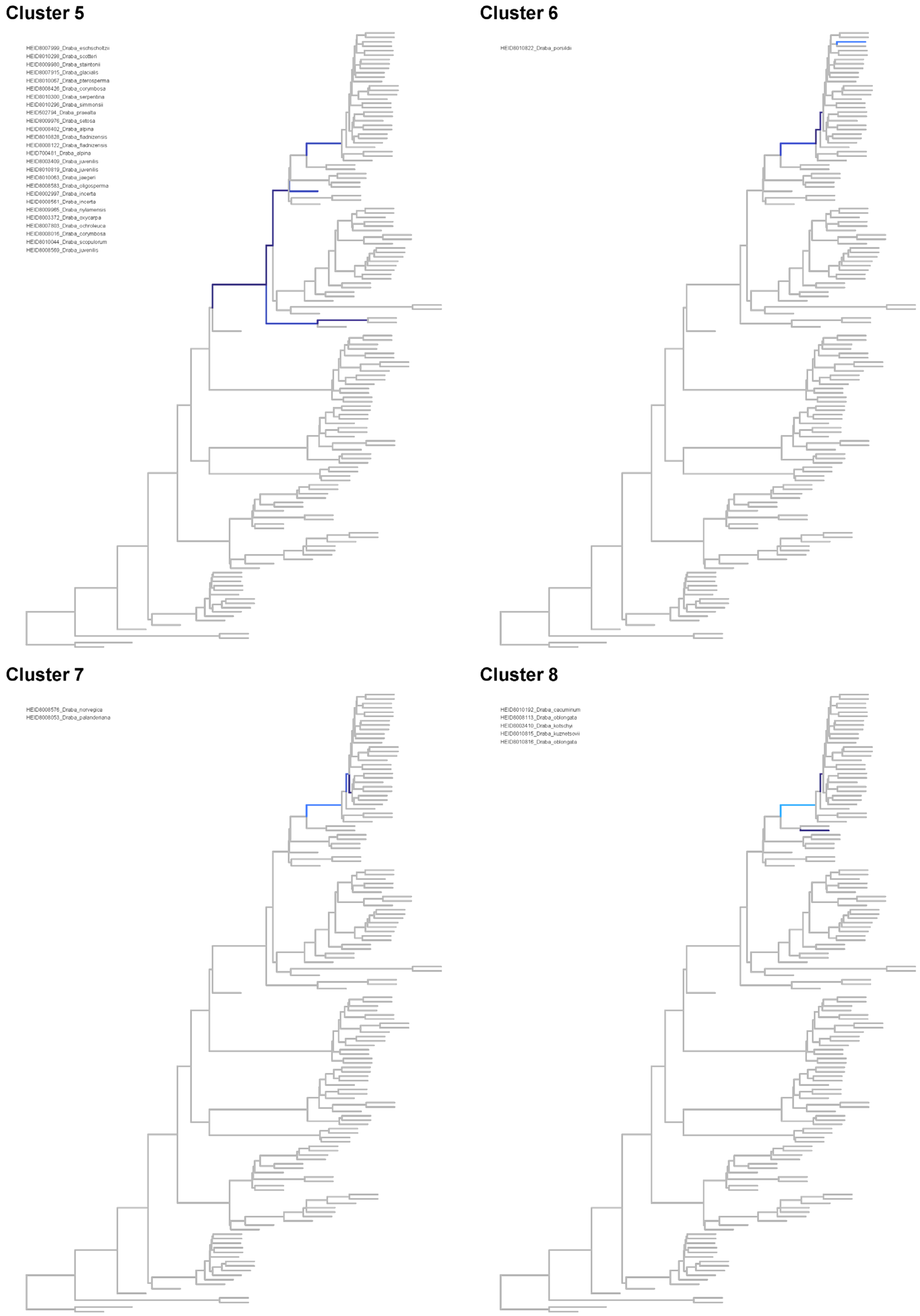

**Figure S20. Dendrograms for paralog placement of clusters 5-8.** Diploid trees with branches colored according to placement proportion are shown; samples assigned to each cluster are given in the top left corner of each subplot. A threshold of 0.05 was chosen to display placement proportion.

**Cluster 9**

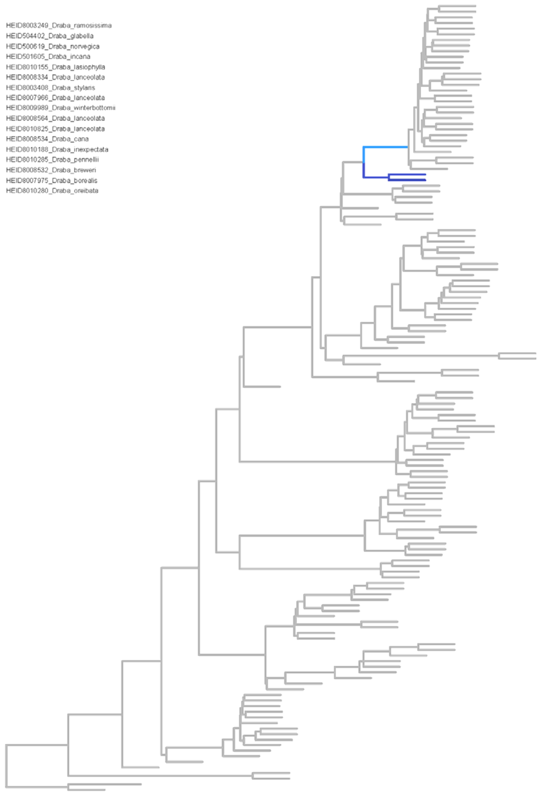

**Cluster 10**

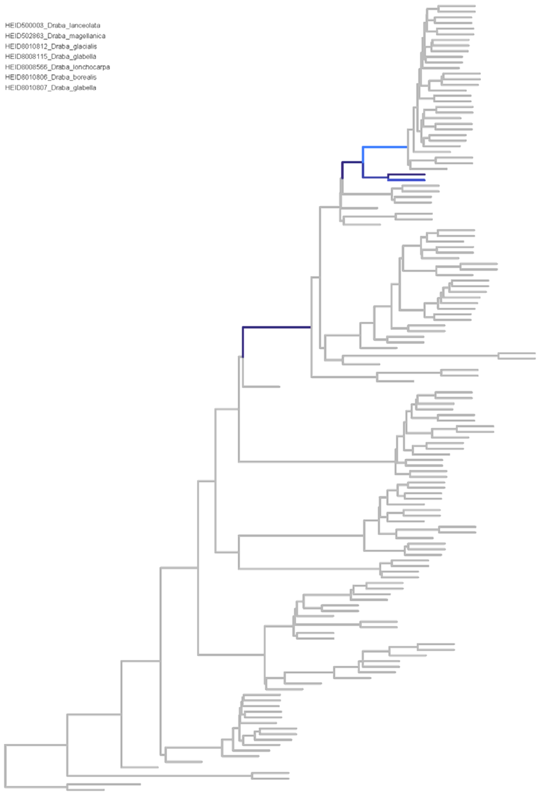

**Cluster 11**

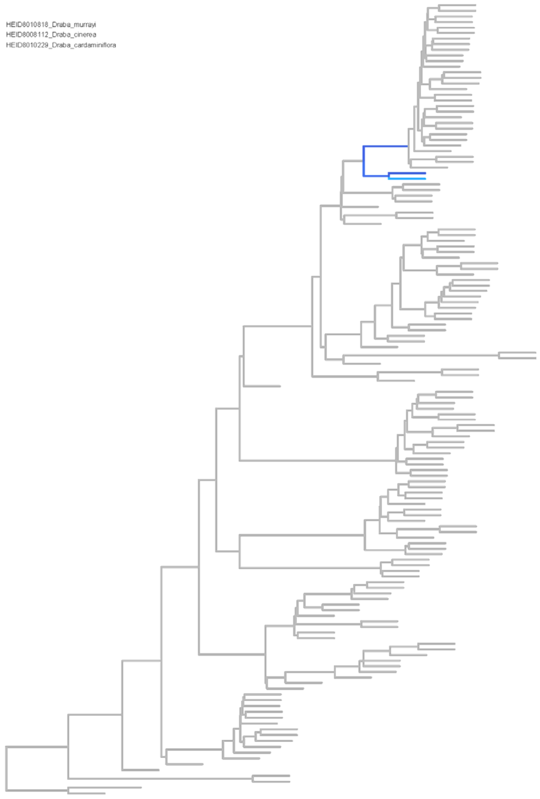

**Cluster 12**

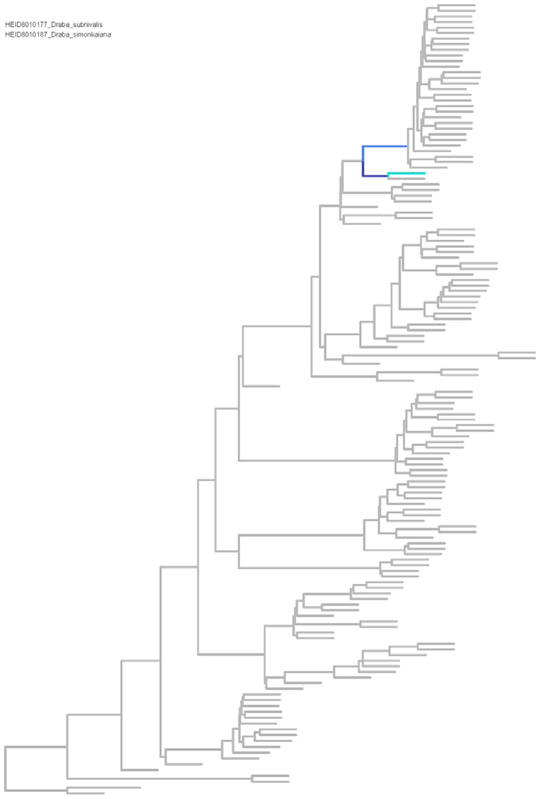

**Figure S21. Dendrograms for paralog placement of clusters 9-12.** Diploid trees with branches colored according to placement proportion are shown; samples assigned to each cluster are given in the top left corner of each subplot. A threshold of 0.05 was chosen to display placement proportion.

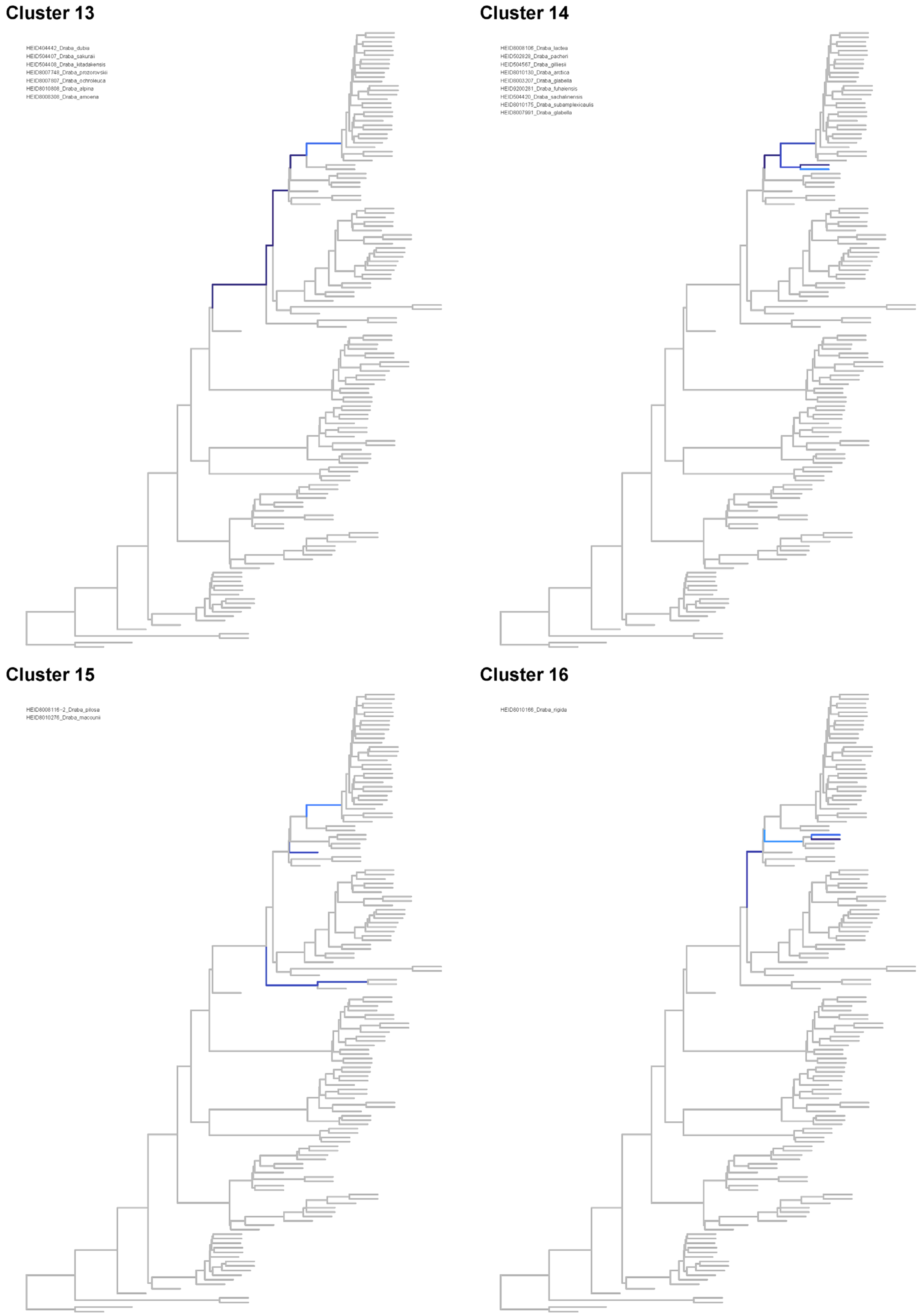

**Figure S22. Dendrograms for paralog placement of clusters 13-16.** Diploid trees with branches colored according to placement proportion are shown; samples assigned to each cluster are given in the top left corner of each subplot. A threshold of 0.05 was chosen to display placement proportion.

**Cluster 17**

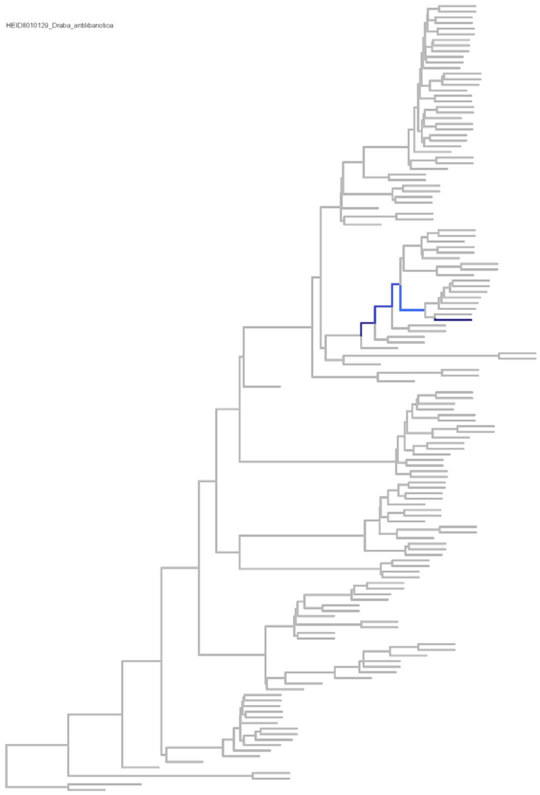

**Cluster 18**

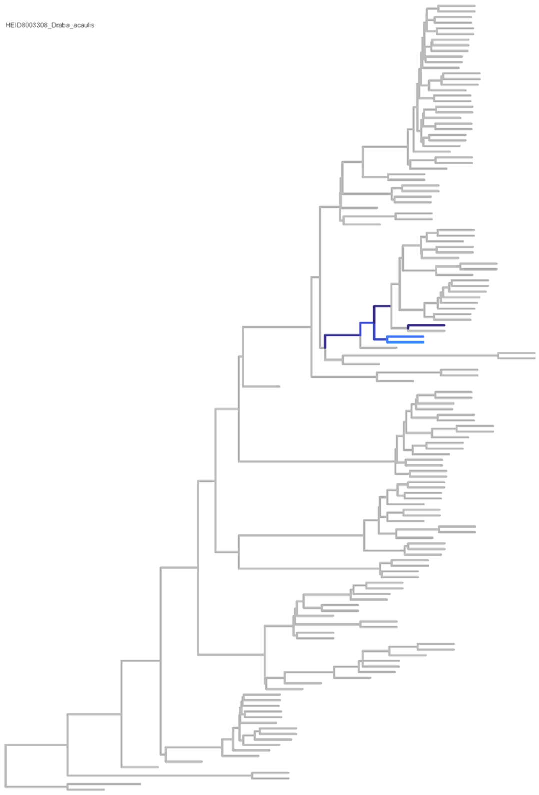

**Cluster 19**

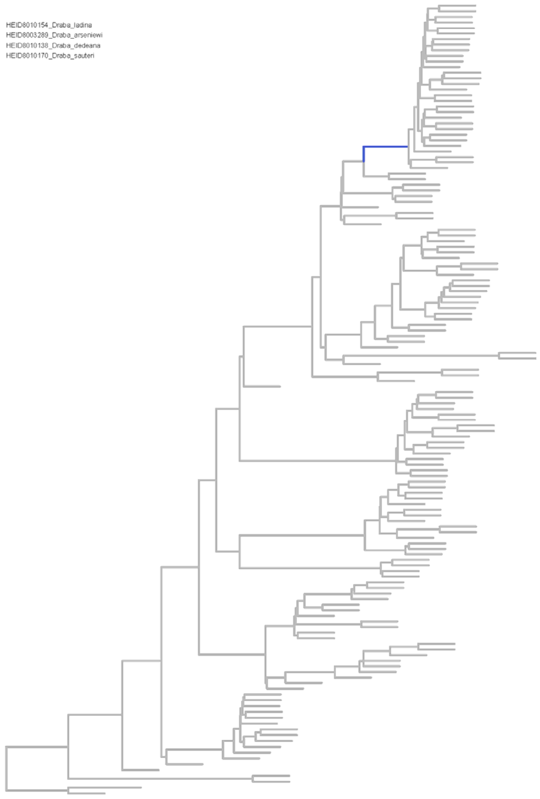

**Cluster 20**

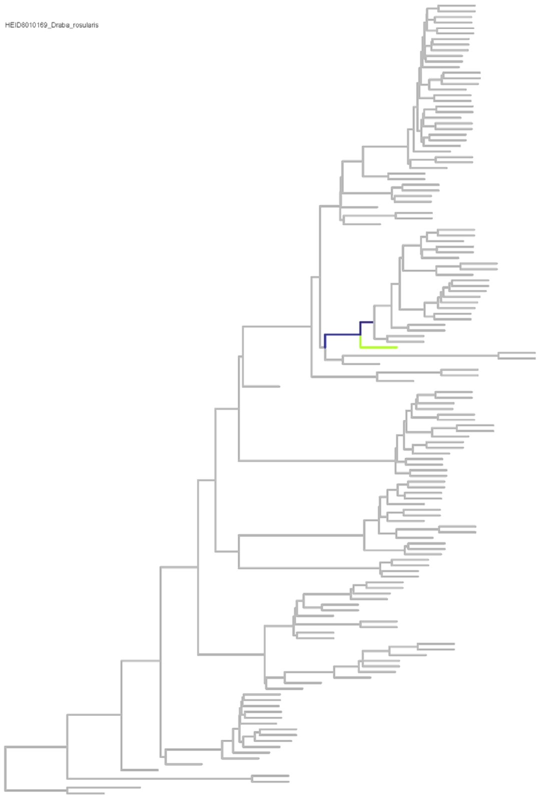

**Figure S23. Dendrograms for paralog placement of clusters 17-20.** Diploid trees with branches colored according to placement proportion are shown; samples assigned to each cluster are given in the top left corner of each subplot. A threshold of 0.05 was chosen to display placement proportion.

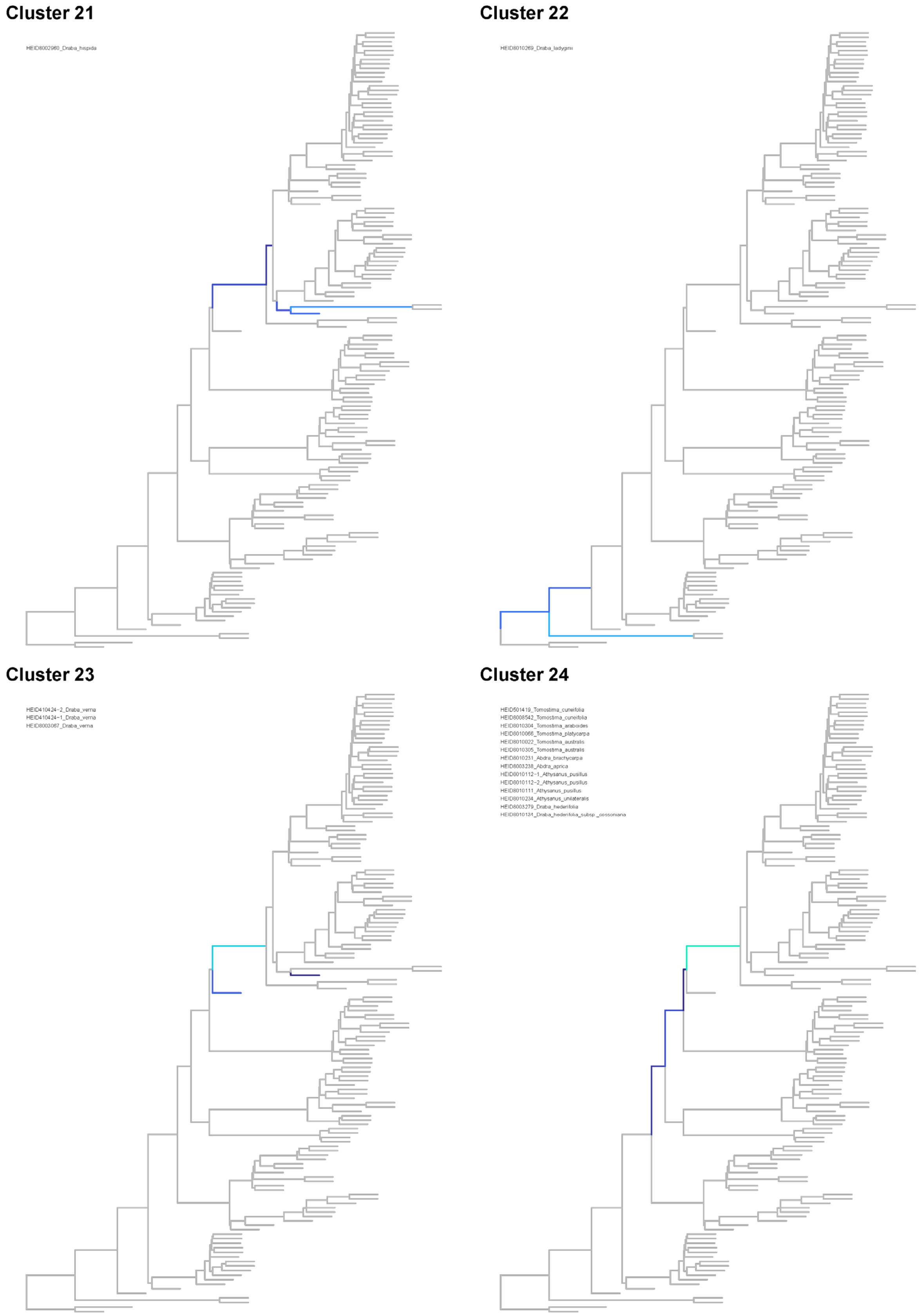

**Figure S24. Dendrograms for paralog placement of clusters 21-24.** Diploid trees with branches colored according to placement proportion are shown; samples assigned to each cluster are given in the top left corner of each subplot. A threshold of 0.05 was chosen to display placement proportion.

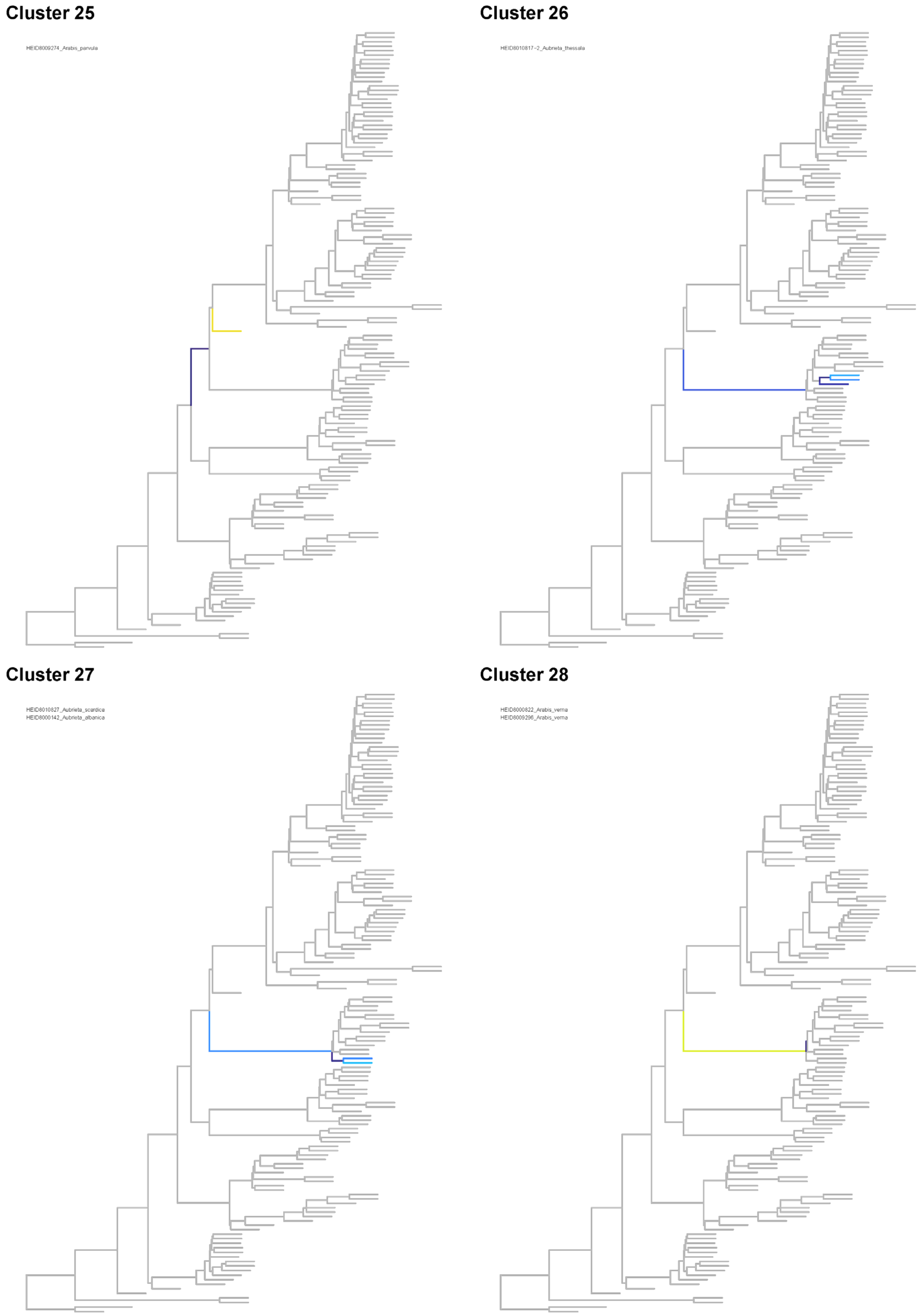

**Figure S25. Dendrograms for paralog placement of clusters 25-28.** Diploid trees with branches colored according to placement proportion are shown; samples assigned to each cluster are given in the top left corner of each subplot. A threshold of 0.05 was chosen to display placement proportion.

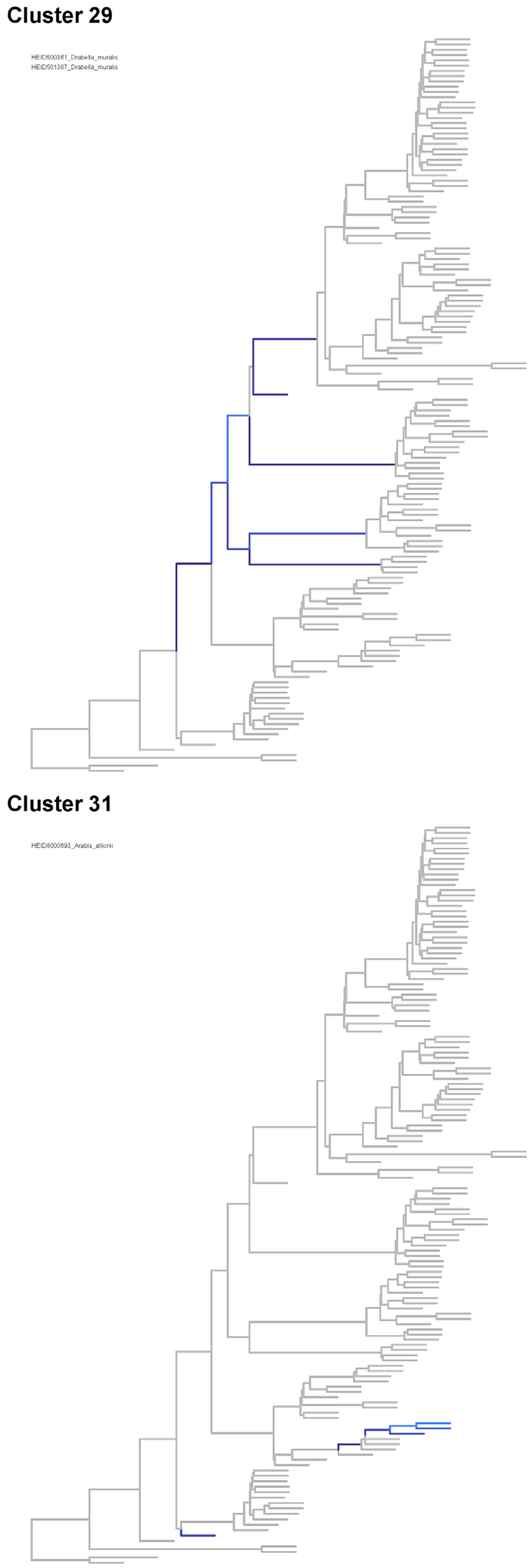

**Figure S26. Dendrograms for paralog placement of clusters 29-32.** Diploid trees with branches colored according to placement proportion are shown; samples assigned to each cluster are given in the top left corner of each subplot. A threshold of 0.05 was chosen to display placement proportion.

Cluster 33

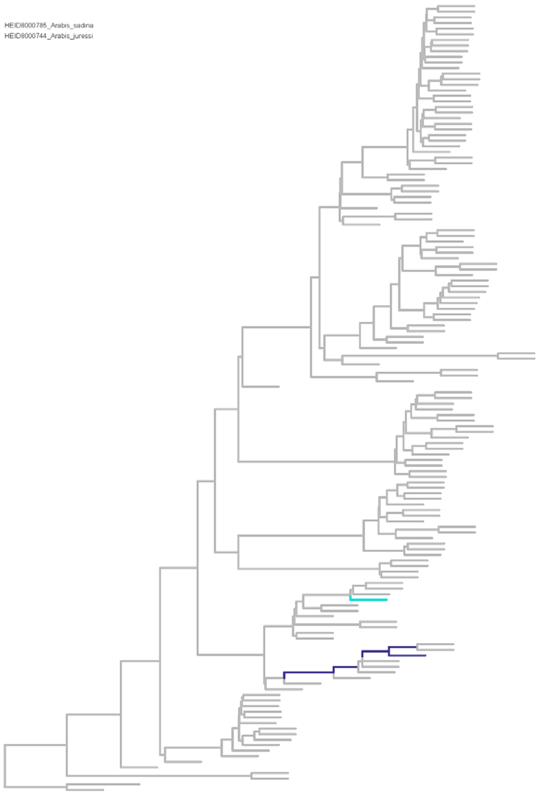

Cluster 34

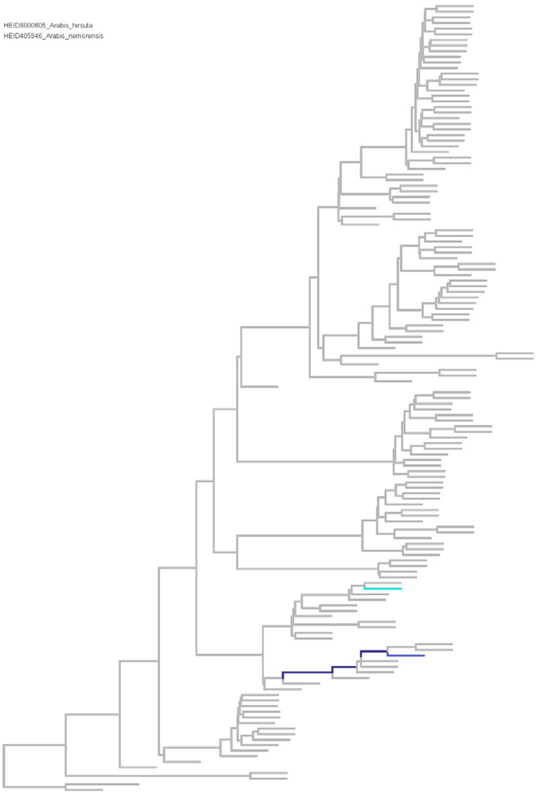

Cluster 35

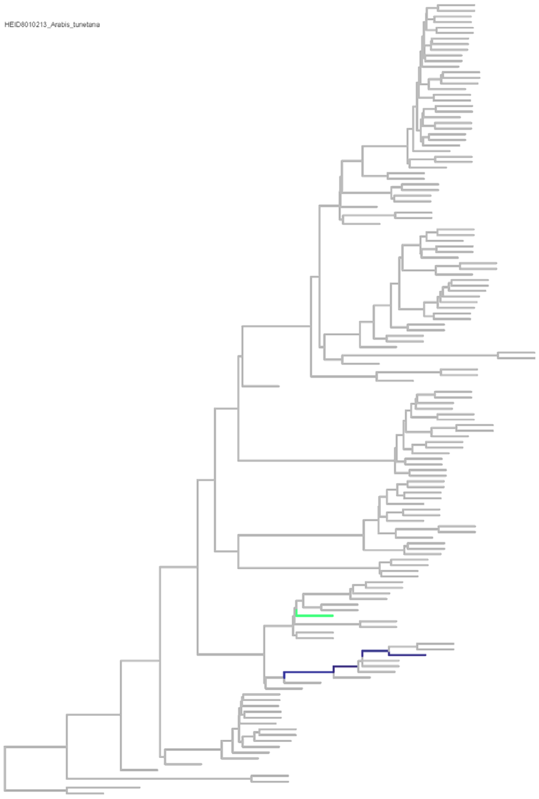

Cluster 36

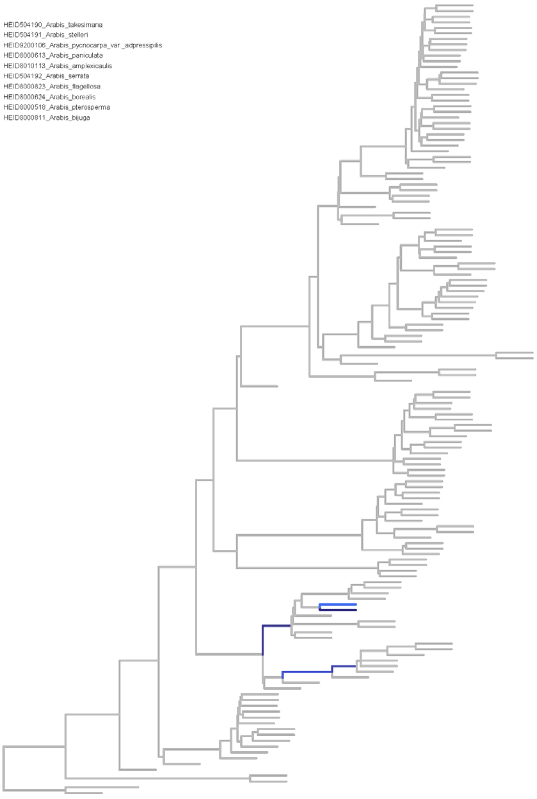

**Figure S27. Dendrograms for paralog placement of clusters 33-36.** Diploid trees with branches colored according to placement proportion are shown; samples assigned to each cluster are given in the top left corner of each subplot. A threshold of 0.05 was chosen to display placement proportion.

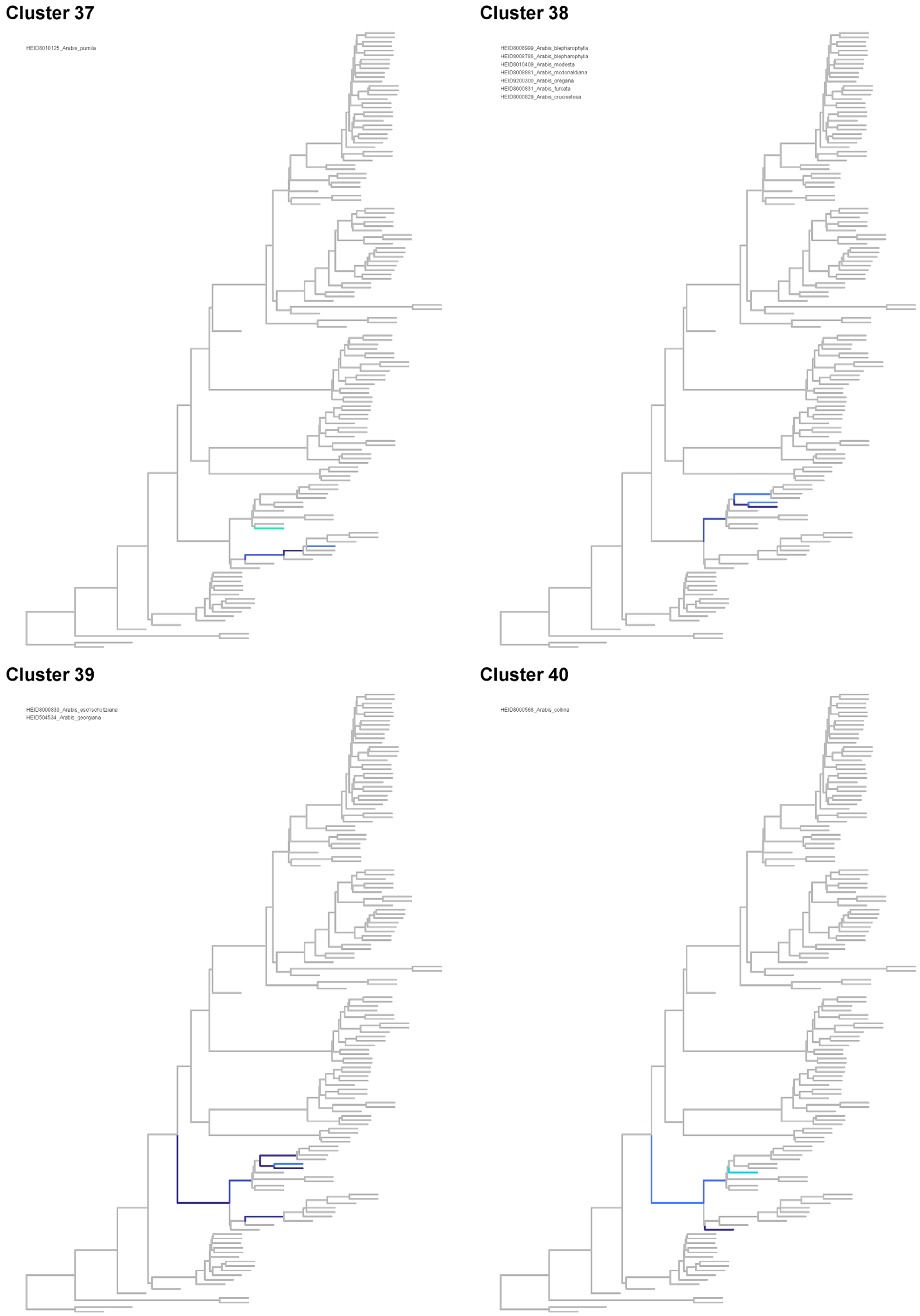

**Figure S28. Dendrograms for paralog placement of clusters 37-40.** Diploid trees with branches colored according to placement proportion are shown; samples assigned to each cluster are given in the top left corner of each subplot. A threshold of 0.05 was chosen to display placement proportion.

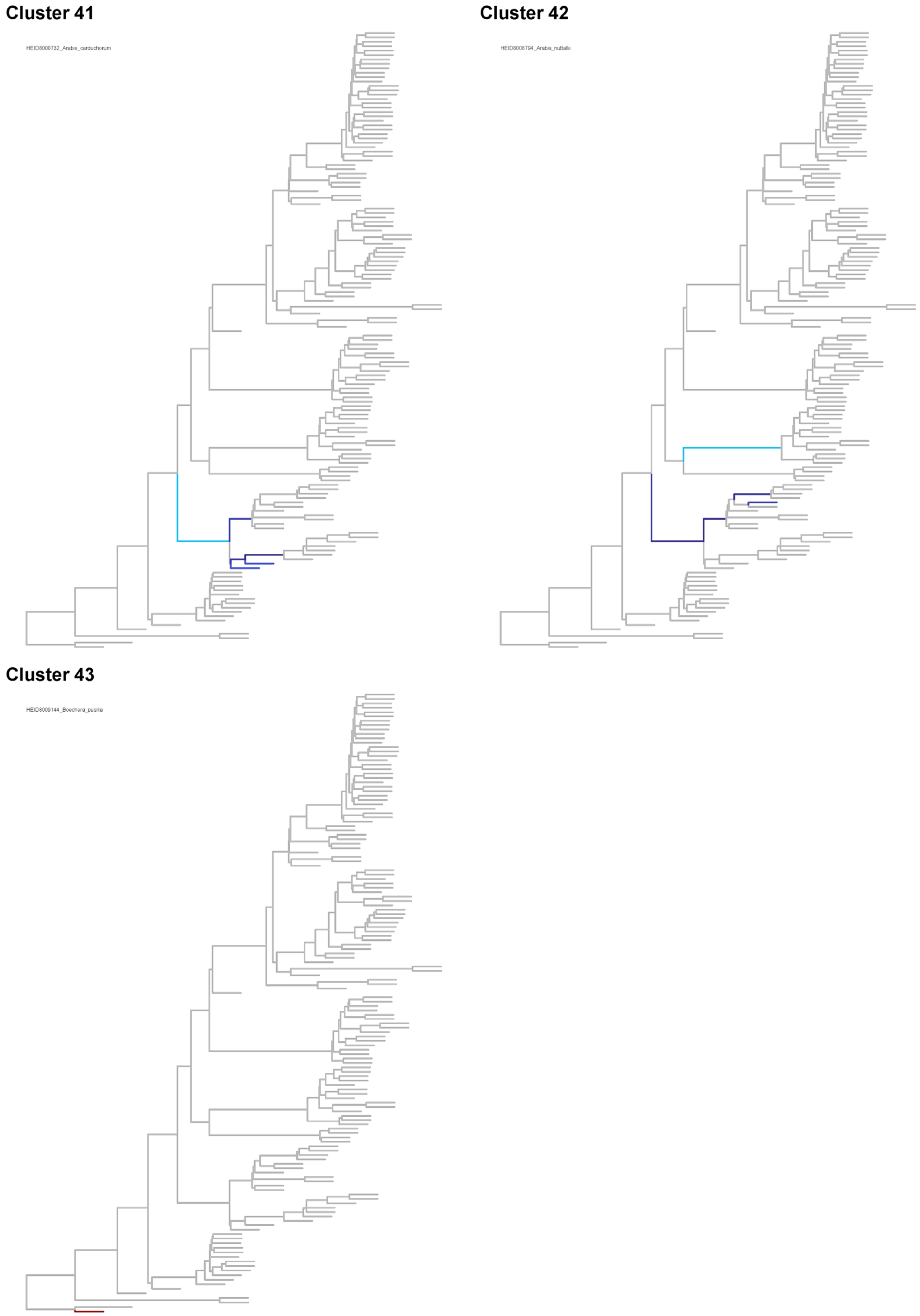

**Figure S29. Dendrograms for paralog placement of clusters 41-43.** Diploid trees with branches colored according to placement proportion are shown; samples assigned to each cluster are given in the top left corner of each subplot. A threshold of 0.05 was chosen to display placement proportion.

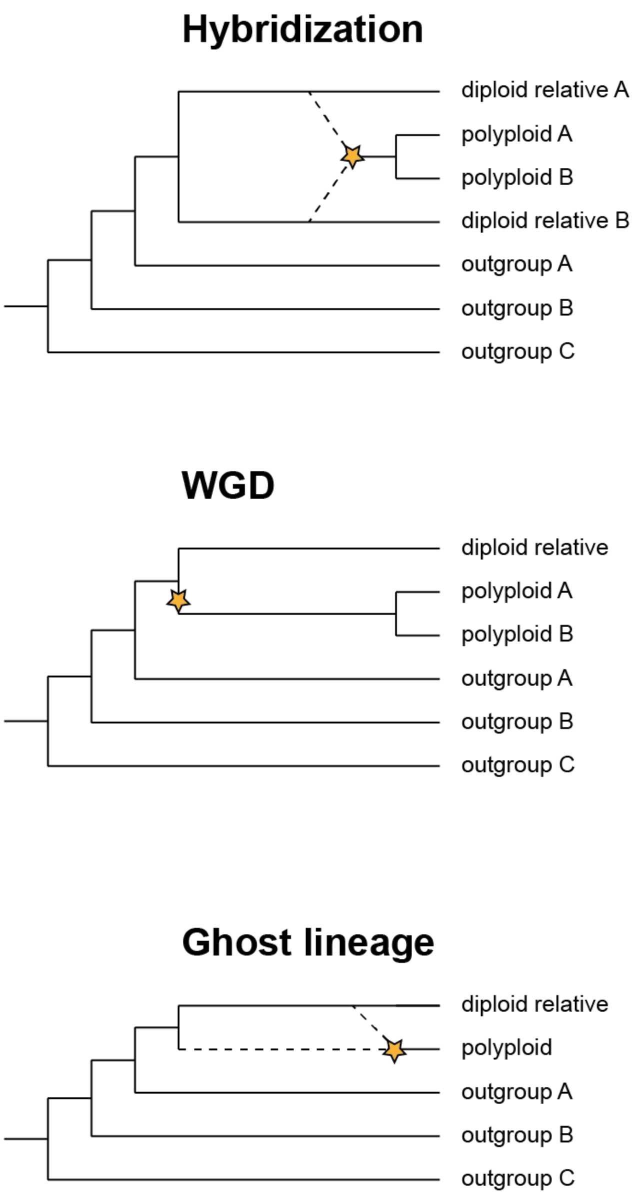

**Figure S30. Three evolutionary scenarios for the emergence of polyploids analyzed with *Ks*.** Hybridization (allopolyploidization) between two diploid lineages leading to the emergence of a polyploid lineage with subsequent lineage diversification, ancient whole-genome duplication (autopolyploidization) at the time of lineage divergence with subsequent lineage diversification, and emergence of a polyploid through recent hybridization of a diploid relative and a now extinct ghost lineage. The orange star symbolizes the emergence of the polyploid taxon/clade.

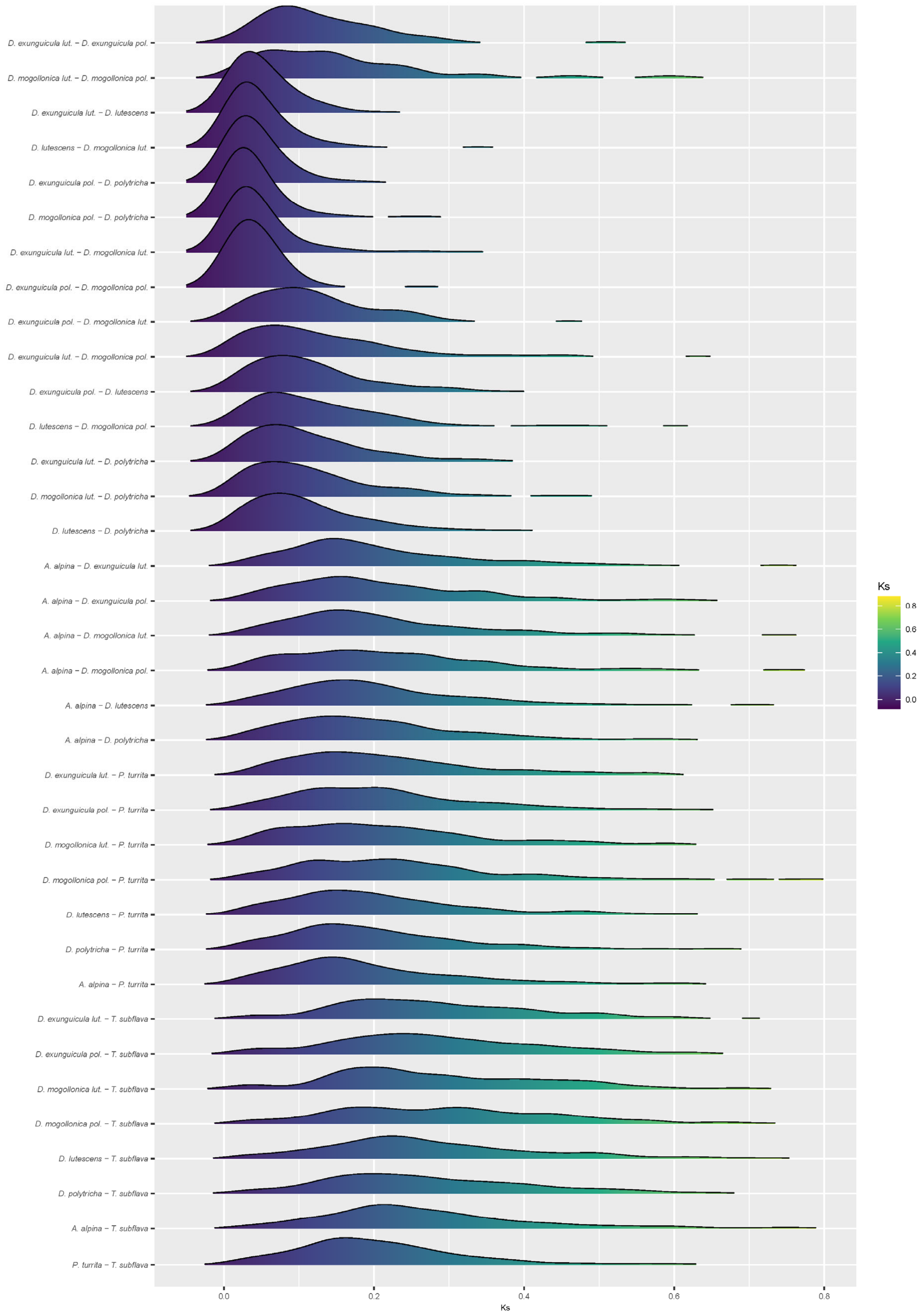

**Figure S31. Synonymous substitution rate distribution in a clade with hybridization.** The distributions of pairwise synonymous substitution rate (*Ks*) of two *Draba* species from cluster 1, *Draba exunguicula* and *Draba mogollonica,* their closest diploid relatives *Draba lutescens* and *Draba polytricha*, as well as *Arabis alpina*, *Pseudoturritis turrita* and *Turritis subflava*, are shown.

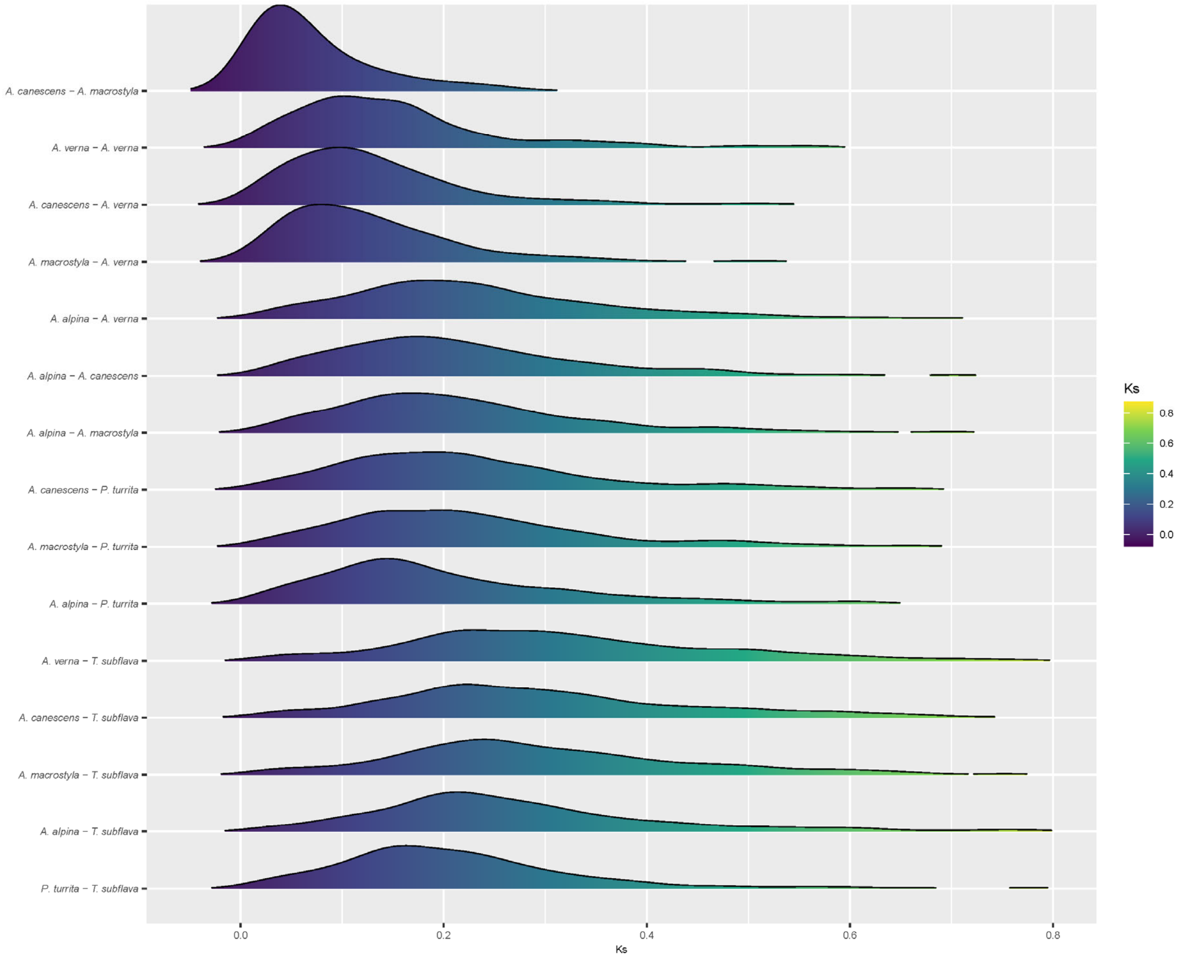

**Figure S32. Synonymous substitution rate distribution in a clade with WGD.** The distributions of pairwise synonymous substitution rate (*Ks*) of the polyploid from cluster 28 (*Arabis verna*) and its closest diploid relatives (*Aubrieta macrostyla* and *Aubrieta canescens*), along with *Arabis alpina*, *Pseudoturritis turrita* and *Turritis subflava*, are shown along with the mean *Ks* of the respective main peaks.

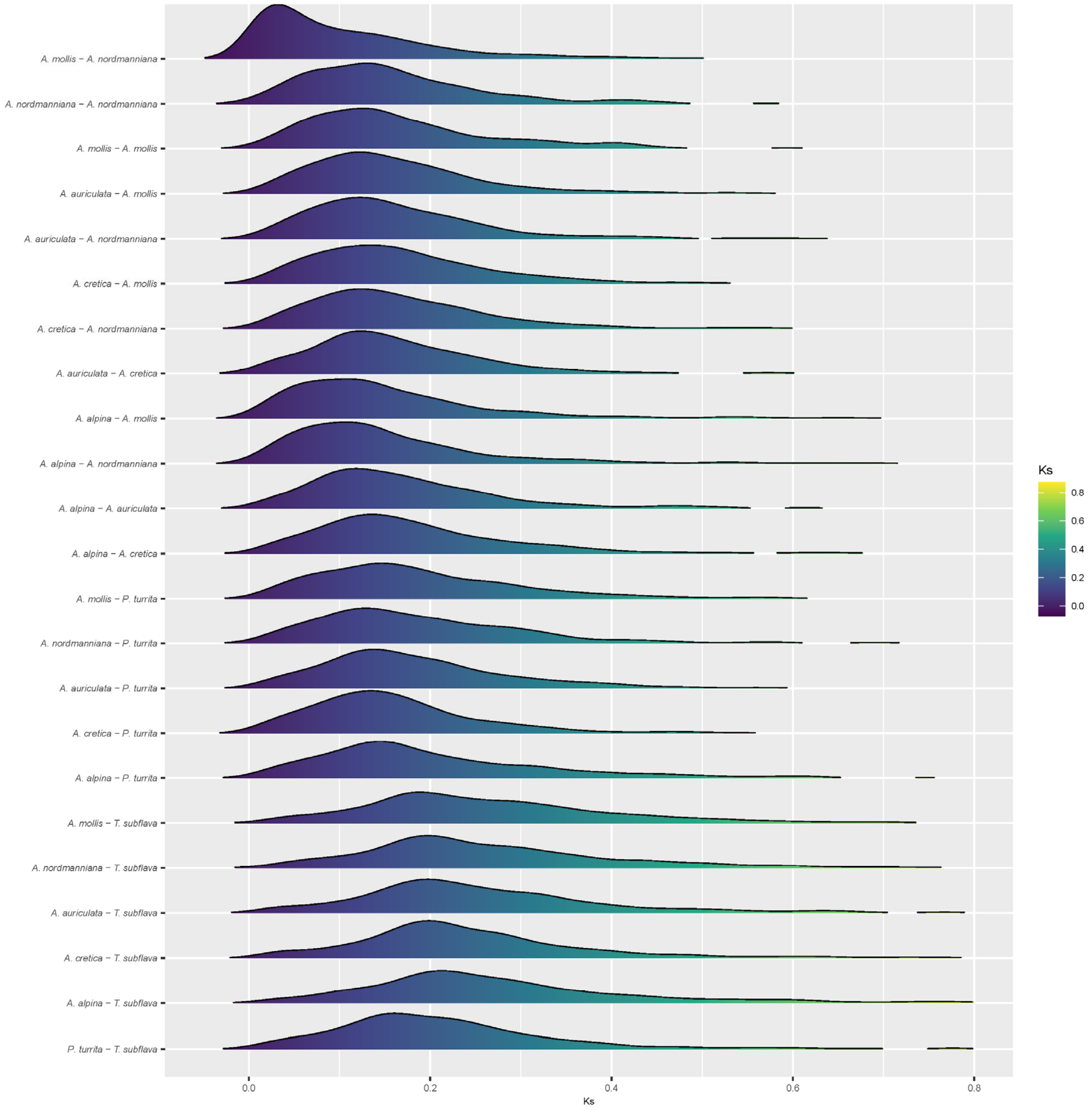

**Figure S33. Synonymous substitution rate distribution in a clade with WGD.** The distributions of pairwise synonymous substitution rate (*Ks*) of two species in cluster 30, *Arabis mollis* and *Arabis nordmanniana*, as well as *Arabis cretica*, *Arabis auriculata*, *Arabis alpina*, *Pseudoturritis turrita* and *Turritis subflava*, are shown along with the mean *Ks* of the respective main peaks.

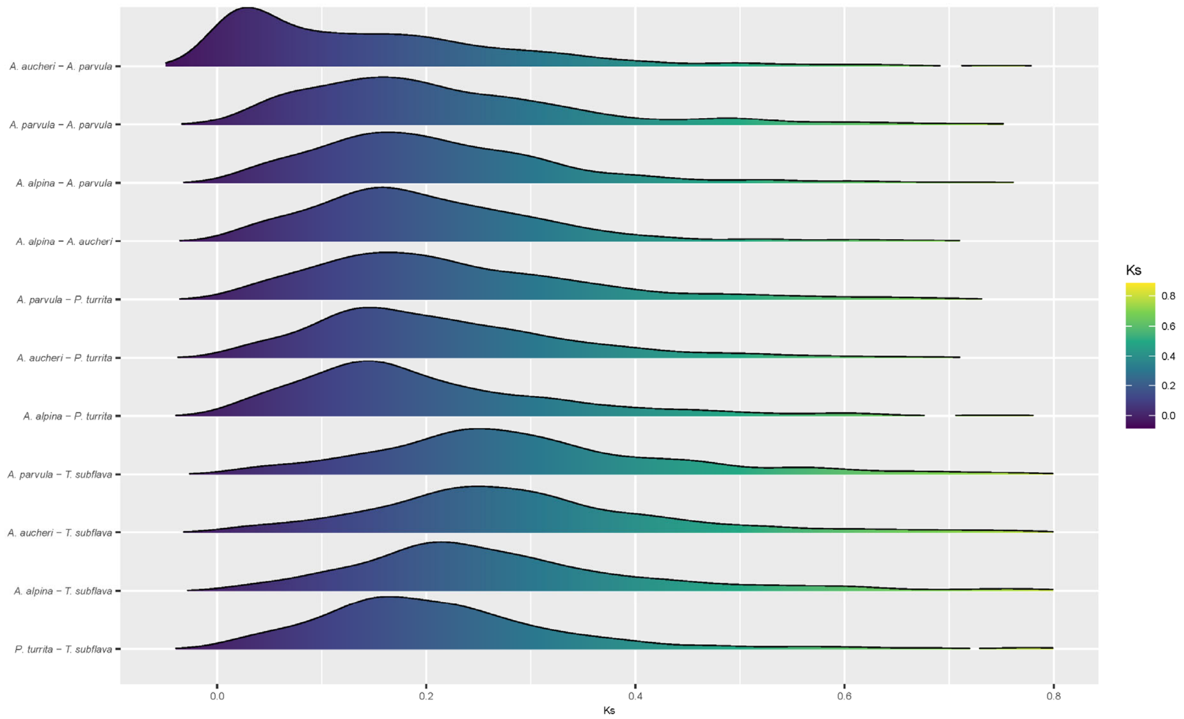

**Figure S34. Synonymous substitution rate distribution in a clade with a ghost lineage.** The distributions of pairwise synonymous substitution rate (*Ks*) of polyploid *Arabis parvula* (cluster 25) and its closest relative *Arabis aucheri*, as well as *Arabis alpina*, *Pseudoturritis turrita* and *Turritis subflava*, are shown.
